# *Zfp189* Mediates Stress Resilience Through a CREB-Regulated Transcriptional Network in Prefrontal Cortex

**DOI:** 10.1101/403733

**Authors:** Zachary S. Lorsch, Peter J. Hamilton, Aarthi Ramakrishnan, Eric M. Parise, William J. Wright, Marine Salery, Ashley Lepack, Philipp Mews, Orna Issler, Andrew McKenzie, Xianxiao Zhou, Lyonna F. Parise, Stephen T Pirpinias, Idelisse Ortiz Torres, Sarah Montgomery, Yong-Hwee Eddie Loh, Benoit Labonté, Andrew Conkey, Ann E. Symonds, Rachael Neve, Gustavo Turecki, Ian Maze, Yan Dong, Bin Zhang, Li Shen, Rosemary C. Bagot, Eric J. Nestler

## Abstract

Stress resilience involves numerous brain-wide transcriptional changes. Determining the organization and orchestration of these transcriptional events may reveal novel antidepressant targets, but this remains unexplored. Here, we characterize the resilient transcriptome with co-expression analysis and identify a single transcriptionally-active uniquely-resilient gene network. *Zfp189*, a previously unstudied zinc finger protein, is the top network key driver and its overexpression in prefrontal cortical (PFC) neurons preferentially activates this network, alters neuronal activity and promotes behavioral resilience. CREB, which binds *Zfp189*, is the top upstream regulator of this network. To probe CREB-*Zfp189* interactions as a network regulatory mechanism, we employ CRISPR-mediated locus-specific transcriptional reprogramming to direct CREB selectively to the *Zfp189* promoter. This single molecular interaction in PFC neurons recapitulates the pro-resilient *Zfp189-*dependent downstream effects on gene network activity, electrophysiology and behavior. These findings reveal an essential role for *Zfp189* and a CREB-*Zfp189* regulatory axis in mediating a central transcriptional network of resilience.

## Introduction

Despite the high prevalence of Major Depressive Disorder (MDD), and the established relationship between major life stressors and MDD, most people who experience stress do not develop depression. This variability in stress susceptibility vs. resilience is likely the result of multiple interrelated factors including genetic differences (CONVERGE, 2015; Hyde et al., 2016), hypothalamic-pituitary-adrenal stress axis function (Herman and Cullinan, 1997) and altered immune reactivity (Hodes et al., 2014). Divergent phenotypic responses to stress are studied in animal models such as chronic social defeat stress (CSDS), where a portion of stressed mice display several MDD-like behaviors (termed ‘susceptible’), while the remainder do not (termed ‘resilient’) (Berton et al., 2006; Krishnan et al., 2007). In an attempt to unravel the neurobiological underpinnings of this resilient state, numerous studies have focused on the molecular (Dias et al., 2014; Taliaz et al., 2011; Vialou et al., 2010; Wilkinson et al., 2011), circuit (Chaudhury et al., 2013; Friedman et al., 2014, 2016; Muir et al., 2018) and epigenetic (Bagot et al., 2014; Elliott et al., 2010; Wilkinson et al., 2009; Zannas and West, 2014) changes that occur uniquely in stress-resistant animals. Importantly, while resilient animals behave similarly to unstressed controls, resilience involves cellular and molecular adaptations that are distinct from those that occur in both unstressed and stress-susceptible animals (Friedman et al., 2014; Krishnan et al., 2007). Accordingly, resilience is not simply the absence of susceptibility, but rather an active homeostatic response to stress. In support of this, RNA-sequencing (RNA-seq) studies report broad transcriptional changes across brain regions, with far more early changes in transcription observed in stress-resilient mice than in stress-susceptible mice (Bagot et al., 2016a; Krishnan et al., 2007). However, the relationship among resilience-associated genes, as well as their mechanistic regulation, has not been determined.

By clustering genes into separate units (modules) based on their coordinated transcriptional regulation, network-based analytical methods such as weighted gene co-expression network analysis (WGCNA) provide a data-driven approach to analyze large transcriptional datasets. WGCNA has been used to define gene networks from human post-mortem brain in numerous complex disorders such as schizophrenia (Maschietto et al., 2015), Parkinson’s disease (Yue et al., 2017), Alzheimer’s disease (Zhang et al., 2013), autism (Parikshak et al., 2013) and MDD (Labonte et al., 2017; Malki et al., 2015). While these studies have successfully utilized *in silico* methods to define relationships among affected genes, they are limited in that the causality of these relationships cannot be probed *in vivo* in humans. Consequently, recent studies from our group have defined stress-susceptibility networks in mouse brain and demonstrated that manipulation of key hub genes within a network drive regulation of the downstream gene network as well as behavioral susceptibility (Bagot et al., 2016a; Labonte et al., 2017). However, the functioning of resilient WGCNA networks has not been explored. Additionally, while these recent studies identify a causal role for key hub genes in network activation, there has been no investigation into how hub genes themselves are regulated, and whether these regulatory mechanisms also control gene network activity. Consequently, there is currently a limited understanding of the higher-order orchestration of coordinated patterns of gene expression, specifically the mechanisms by which transcriptional networks are activated and silenced. Given the considerable overlap in transcriptional (Bagot et al., 2016b) and epigenetic (Wilkinson et al., 2009) changes induced in the resilient brain and those induced by antidepressant treatment, detailed knowledge of the organization and regulation of the resilient transcriptome could provide novel therapeutic targets for MDD.

In the present study, we employ a stepwise approach to define WGCNA networks unique to the stress resilient phenotype and explore their mechanistic regulation *in vivo*. Because the resilience phenotype is distinct from both stress susceptibility and control conditions, we focus on transcriptional networks that are uniquely configured in the resilient state. To understand the precise mechanisms of network regulation, we explore both key driver genes within networks as well as their upstream regulators and define the specific molecular interactions between the two. Using this approach, we discover *Zfp189*, a putative zinc-finger transcription factor about which virtually nothing is known, as the strongest hub gene in the most highly ranked transcriptionally-active resilient-specific gene module, and find that CREB is the strongest predicted upstream regulator of genes within this module. We show that *Zfp189* overexpression in prefrontal cortex (PFC), the brain region implicated most strongly in our WGCNA analysis, drives altered expression of module genes and promotes stress resilience. Furthermore, we establish a casual interaction between CREB and *Zfp189*, which we recreate *in vivo* via a novel application of CRISPR technology to direct CREB binding specifically to the *Zfp189* gene promoter in PFC neurons, showing that this single regulatory interaction controls network-wide gene expression, PFC neuron activity, and behavioral resilience. These findings demonstrate a key role for *Zfp189*, its downstream gene network activity, and its upstream regulatory control, in driving the molecular events central for stress resilience.

## Results

### *Zfp189* Regulates a Resilient-Specific Transcriptional Network

To identify gene modules within the broad gene expression changes of stress resilience, we applied WGCNA to our recently published RNA-seq data set of mice resilient to CSDS (Figure 1A) (Bagot et al., 2016a). This approach reveals 30 modules (color names are arbitrary) of gene expression present across four limbic brain regions (Figure 1B). To confirm the validity of this approach, we looked for the presence of known biological relationships within our modules via protein-protein interactions (PPI) and cell-type specific genes, finding several modules enriched for each (Figure S1A-B). To determine the relevance of these networks to human MDD, we examined whether the CSDS-associated resilience modules are preserved in RNA-seq data from post-mortem MDD patients and matched controls (Labonte et al., 2017). 56.6% of our modules are preserved in human brain, but—consistent with a role in resilience—more modules (11 vs. 2) show greater preservation in control conditions than in MDD (Figure S1C-F).

**Figure 1.**
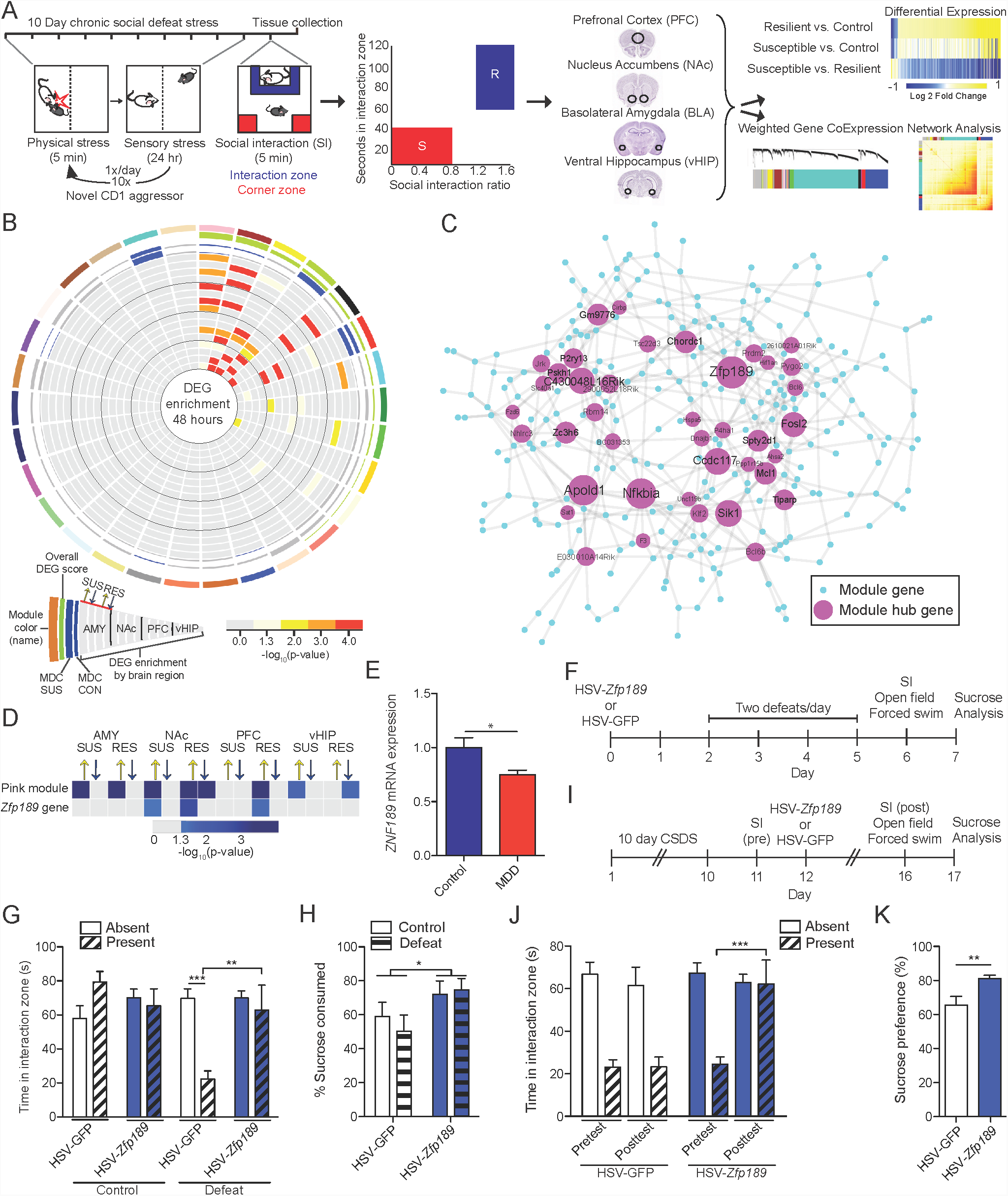
Identification of the Resilient-Specific Pink Module and its Pro-Resilient Top Key Driver *Zfp189*. (A) Overview of chronic social defeat stress (CSDS) protocol, social interaction test (SI) phenotyping, brain dissections, and RNA-sequencing analysis of four brain regions, prefrontal cortex (PFC), nucleus accumbens (Nac), basolateral amygdala (BLA) and ventral hippocampus (vHIP), used to identify resilient-specific transcriptional networks. (B) Resilient modules (colored bars) identified by weighted gene co-expression network analysis (WGCNA). Modules, named with an arbitrary color (outer most ring) are ranked clockwise by overall differentially expressed gene (DEG) enrichment (p < 0.05, FC > 1.3). Phenotype specificity as determined by resilient module differential connectivity (MDC) in susceptible and control networks and DEG enrichment is displayed internally. Presence of MDC color denotes statistical significance (FDR q < 0.05). DEG enrichments are scaled by -log_10_(p-value) with only significant (p < 0.05) enrichments featured in color. The pink module (top) is the only module that shows DEGs across brain regions and MDC when compared to both susceptible and control mice. (C) Network structure of the pink module. Key drivers are featured and scaled in size according to number of connections in the network. *Zfp189* is the top key driver gene. (D) Correspondence between differential expression for *Zfp189* and DEG enrichment for the pink module across phenotypes and brain areas. DEG enrichments are scaled by -log_10_(p-value) with only significant (p < 0.05) enrichments featured in color. In PFC, both *Zfp189* and the pink module are only affected in animals resilient to CSDS (both upregulated). (E) mRNA of *ZNF189*, the human ortholog of *Zfp189*, is reduced in post-mortem PFC from major depressive disorder (MDD) patients (n = 17-22). (F) Experimental timeline to characterize behavioral effects of virally overexpressing *Zfp189* in PFC prior to CSDS. (G) Pro-resilient behavioral effects of *Zfp189* in SI. Mice injected intra-PFC with HSV-*Zfp189* and exposed to CSDS spend more time in the interaction zone when a target mouse present than defeated HSV-GFP mice (n= 9-13). (H) Mice overexpressing *Zfp189* in PFC have an elevated preference for sucrose relative to HSV-GFP mice (n = 9-14). (I) Experimental timeline to determine behavioral effects of overexpressing *Zfp189* in PFC in CSDS-susceptible mice. (J) *Zfp189* reverses depression-like social withdrawal in susceptible mice. Susceptible mice injected intra-PFC with HSV-*Zfp189* spend more time in the interaction zone when the target mouse is present in the post-injection posttest than the pre-injection pretest, but HSV-GFP injection does not change behavior (n = 11-13). (K) Previously susceptible mice injected with HSV-*Zfp189* have a higher sucrose preference than previously susceptible mice injected with HSV-GFP (n = 11-12). *p < 0.05, **p < 0.01, ***p < 0.001. Bar graphs show mean ± SEM. See also Figures S1-S2 and Tables S1-S2.

We next sought to identify the transcriptional relationships that are unique to the resilient phenotype. While all resilient modules by definition consist of genes co-expressed in the resilient brain, it is likely that some of these networks are preserved in susceptibility, control or both conditions. To identify resilient-specific modules, we performed module differential connectivity (MDC) analysis (Zhang et al., 2013) to compare resilient modules to networks generated from susceptible mice and unstressed controls (Bagot et al., 2016a). This analysis determines whether the relationship between genes within a resilient module is preserved in the different conditions. In this context, modules can gain connectivity in which expression of genes becomes more correlated, lose connectivity in which expression of genes becomes less correlated, or show no change. While 22 of 30 resilient modules show no change in connectivity, three modules show significant MDC with only the susceptible phenotype, one module shows significant MDC with only the control phenotype, and four modules show significant MDC with both phenotypes (Figure 1B). We also tested the degree to which each resilient module contains genes that are differentially expressed in the four brain regions studied after CSDS. Of our 30 resilient modules, 12 are enriched for differentially expressed genes (DEGs) 48 hours post-stress (Figure 1B).

We next combined MDC and DEG analyses to identify resilient-specific and transcriptionally-active networks. Of the 12 modules enriched for DEGs, only two, the pink and brown modules, were enriched in all brain regions studied (Figure 1B). While brown module connectivity is distinct from susceptible (MDC = 0.9), pink module connectivity is distinct from both susceptible (MDC = 0.89) and control (MDC = 0.91). Therefore, only the pink module is entirely resilient-specific and differentially expressed across all brain regions studied. In further support of the importance of this network, the resilient-specific pink module is enriched for both PPIs and cell-type specific genes (Figure S1) and more strongly preserved in human controls (p = 1.07 x 10^−17^) than in human MDD (p = 1.14 x 10^−15^) (Figure S1F).

To determine the structure of the pink module, we reconstructed the network based on individual gene-gene correlations using the Algorithm for Reconstruction of Accurate Cellular Networks (Margolin et al., 2006) and predicted gene regulators with Key Driver Analysis (Zhang and Zhu, 2013) (Figure 1C). Of 281 genes in the network, 40 meet criteria for key driver genes (Table S1). The top key driver gene is *Zfp189*, which contains 52 defined connections and is tightly integrated in the core network with eight direct connections to other key driver genes (Figure S2A-B). To determine the brain regions in which these key driver genes regulate the pink module, we compared individual differential expression patterns of *Zfp189* and the other top 10 key driver genes to module DEGs across brain regions (Figure 1D; Figure S2C). Of these 10 key drivers only *Zfp189* and *Nfkbia* are differentially expressed, and only *Zfp189* recapitulates the brain-region specific DEG enrichment profile of the pink module (both are upregulated in PFC). As the PFC is robustly implicated in stress resilience (Bagot et al., 2016a), with optogenetic stimulation of this region increasing resilient behavior (Bagot et al., 2015; Covington et al., 2010), this transcriptional overlap is in line with a functional role for *Zfp189* in driving stress resilience through the resilient-unique pink module. However, other than its deduced protein structure (Odeberg et al., 1998), virtually nothing is known about *Zfp189*.

### *Zfp189* in PFC Exerts Pro-Resilient and Antidepressant-Like Actions

To examine the relevance of *Zfp189* to human resilience, we first performed qPCR to evaluate expression of *ZNF189*, the human ortholog of *Zfp189*, in PFC of post-mortem brains of humans with MDD (Table S2). Consistent with its deduced role in promoting resilience to CSDS, *ZNF189* mRNA is reduced in human MDD (U = 110.0, p = 0.0303, n = 17-22; Figure 1E).

Having shown that *Zfp189* is increased in PFC of animals resilient to CSDS and decreased in PFC in human MDD, we next tested the behavioral effects of *Zfp189* in this brain region. By examining RNA expression across cortical cell types in a single-cell RNA-seq study in mouse (Zeisel et al., 2015), we found that *Zfp189* is highly enriched in neurons. Therefore, we used herpes simplex virus (HSV) vectors, which target neurons selectively, to overexpress *Zfp189* plus GFP or GFP alone in PFC neurons and exposed mice to an accelerated social defeat paradigm (Figure 1F; Figure S2D). In the social interaction test (SI), defeated mice overexpressing *Zfp189* in PFC spend more time interacting with a social target than defeated mice with GFP alone, but this is not the case when the social interaction zone is empty or in stress-naïve mice (F_1,40_ = 8.501, p = 0.006, n = 9-13, Bonferroni post hoc p < 0.01; Figure 1G). Consequently, defeated HSV-*Zfp189* mice are resilient to this behavioral effect of social defeat. To determine whether this pro-resilient effect of *Zfp189* extends to other measures of depression- and anxiety-like behavior, we analyzed behavior in the open field test (OFT), forced swim test (FST), and sucrose preference test. In OFT, defeated HSV-GFP mice spend less time in the center of the open field than unstressed HSV-GFP mice (*χ*^*2*^(3) = 10.903, p = 0.012, n = 8-14, Mann Whitney post hoc p = 0.031; Figure S2E), but HSV-*Zfp189* mice do not show this stress-induced anxiety-like effect, although unstressed HSV-*Zfp189* mice show reduced time in center compared to HSV-GFP controls. Notably, locomotor differences do not explain these behavioral effects as there is an effect of CSDS, but not virus, on overall movement (F_1,41_ = 30.26, p < 0.001, n = 9-14; Figure S2F). While there are no FST differences (*χ*^*2*^(3) = 0.564, p = 0.905, n = 9-14; Figure S2G), HSV-Zfp189 mice consume significantly more sucrose than HSV-GFP mice in the sucrose preference test (F_1,41_ = 5.102, p = 0.029, n = 9-14, Figure 1H). Together, these results are consistent with reduced depression-like behavior in mice upon increased neuronal expression of *Zfp189* in PFC.

Since antidepressant treatment is only initiated after a diagnosis of MDD, molecular targets that reverse depression-like behavior have higher therapeutic potential than targets which prophylactically prevent susceptibility. We therefore explored the antidepressant potential of *Zfp189* by generating a cohort of CSDS susceptible mice and overexpressing *Zfp189* plus GFP or GFP alone in PFC to determine whether depression-like behavior could be reversed (Figure 1I). In SI, only *Zfp189* overexpressing mice spend more time in the interaction zone when a target mouse is present in the posttest (after virus was injected) than the pretest (before virus) (interaction F_1,23_ = 5.634, p = 0.026, n = 11-13, Bonferroni post hoc p < 0.001; Figure 1J). HSV-*Zfp189* mice also display a higher sucrose preference than HSV-GFP mice (U = 29.0, p = 0.025, n = 11-12; Figure 1K). As such, *Zfp189* overexpression is sufficient to reverse the symptoms of susceptibility. *Zfp189* overexpression in susceptible mice has no effect in OFT (t = 0.049, p = 0.962, n = 12-13), locomotion testing (t = 0.077, p = 0.450, n = 12-13), or FST (t = 0.142, p = 0.888, n = 12-13) (Figure S2H-J). Thus, *Zfp189* in PFC exerts both pro-resilient and antidepressant-like actions.

To directly test the downstream effects of *Zfp189* on gene expression, we micro-dissected virally-infected tissue from previously susceptible mice that remained susceptible (HSV-GFP) or became resilient (HSV-*Zfp189)* in the SI posttest and performed RNA-seq. Overexpression of *Zfp189* in PFC induces broad changes, with upregulation of 2460 genes and downregulation of 1709 genes (log_2_FC > |0.2|, p < 0.05). Within the pink module, 33.1% (93) of genes are upregulated, including 47.5% (19) of key driver genes (Figure 2A). Many of these genes are tightly integrated in the network structure, including *Nfkbia* and *Apold1*, the two top key driver genes after *Zfp189*. Downregulation was less prominent: 7.1% (20) of genes overall and only 5.0% (2) of hub genes. To determine whether this overlap is statistically significant, we performed a logistic regression-based enrichment analysis to map *Zfp189-*dependent transcriptional changes to all resilient modules. In this network-wide analysis, *Zfp189* overexpression in PFC downregulates 4 and upregulates 12 resilient modules (Figure 2B). In support of our proposed mechanism involving *Zfp189* inducing behavioral resilience through transcriptional regulation of the pink module, the pink module is significantly upregulated (p = 4.45 x 10^−13^). *Zfp189* overexpression also induces the pink module in unstressed mice, although to a lesser extent than with concomitant stress (p = 2.63 x 10^−5^; Figure S3A-B).

**Figure 2.**
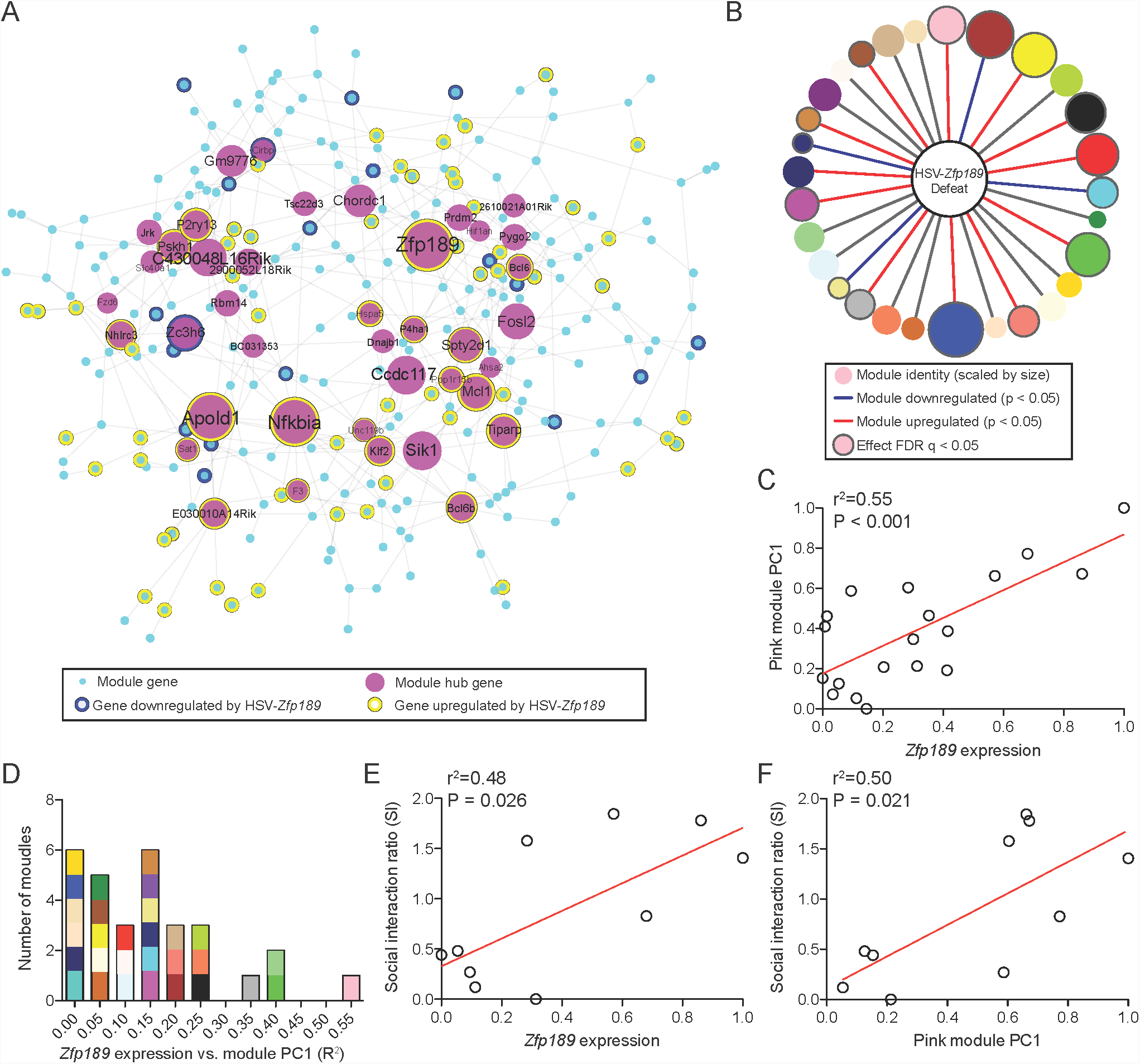
Antidepressant-Like Effects of *Zfp189* Associate With Pink Module Expression Changes. (A) Pink module genes are differentially expressed (p < 0.05, log_2_FC > |0.2|) in PFC following reversal of susceptibility with HSV-*Zfp189*. (B) Module-wide enrichment for HSV-*Zfp189* overexpression in PFC in previously susceptible mice. (C) Variations in *Zfp189* predict the first principal component of pink module expression (n = 19). (D) Of all resilient modules, *Zfp189* shows the strongest regulation of the pink module. (E-F) Positive relationship between resilient behavior and (E) *Zfp189* levels in PFC of each mouse and (F) pink module expression (n = 10). See also Figure S3.

We next tested whether the broad transcriptional effects of *Zfp189* overexpression in PFC (Figure S3C-D) are stronger for the pink module than for other resilient networks. To do this, we performed principal component analysis to determine module expression across HSV-*Zfp189* and HSV-GFP samples in both unstressed and defeated groups. Consistent with a role for *Zfp189* in driving pink module expression, levels of *Zfp189* expression in each PFC sample significantly explain variation in pink module expression as measured by the first principal component (PC1) (R^2^ = 0.55, p < 0.001; Figure 2C). This recapitulates the finding based from our WGCNA analysis (Figure 1C) in an independent dataset. However, while WGCNA is undirected, and because variance in *Zfp189* in this context is virally-induced, these data establish a causal link between *Zfp189* expression and regulation of the pink module. To determine whether this is the case for all resilient modules, we repeated this analysis for all networks. Remarkably, *Zfp189-*induced expression changes drive expression of the pink module more than any other network by far (Figure 2D). We further examined the relationship between variation in *Zfp189* expression, resilient behavior, and pink module expression, and found that all three are related (R^2^ = 0.48, p = 0.026 and R^2^ = 0.50, p = 0.021 respectively; Figure 2E-F). Together, these data validate our bioinformatics predictions and indicate that the pro-resilience effects of *Zfp189* are preferentially associated with pink module gene expression.

### CREB is an Upstream Regulator of the Pink Module

Having demonstrated a causal role for the top module hub gene *Zfp189* in activating the resilient-specific pink module, we next sought to investigate external mechanisms of module regulation. First, we used HOMER (Heinz et al., 2010) to evaluate pink module genes for overrepresentation of known binding motifs *in silico*. Both activating transcription factor 1 (ATF1) and cyclic AMP (cAMP) response element (CRE) binding protein (CREB) are predicted upstream regulators of pink module genes, with consensus sites in 25.3% and 19.7% of pink module genes, respectively (FDR q = 0.027 and FDR q = 0.035; Figure 3A and Table S3). Notably, ATF1 and CREB are closely-related members of a subfamily of leucine zipper transcription factors that bind CRE sites to regulate gene expression. Indeed, both ATF1 and CREB predictions are predicated on a near-identical CRE-containing consensus sequence, indicating that the CRE site is overrepresented in the pink module. To determine the specificity of this prediction to the pink module, we examined other resilient modules. Statistical enrichment of any binding site was observed in only four of the 29 other networks (FDR q < 0.05; Table S3), and none implicated the CRE sequence. Both ATF1 and CREB play important roles in cell survival (Bleckmann et al., 2002) and are ubiquitously activated by extracellular signals (Shaywitz and Greenberg, 1999), yet CREB has also been implicated extensively in both human MDD (Juhasz et al., 2011; Xiao et al., 2017) and animal models of depression (Carlezon et al., 2005; Chen et al., 2001; Covington et al., 2011; Wilkinson et al., 2009), while only non-ATF1 members of the ATF family have been implicated in stress responsivity (Green et al., 2008).

**Figure 3.**
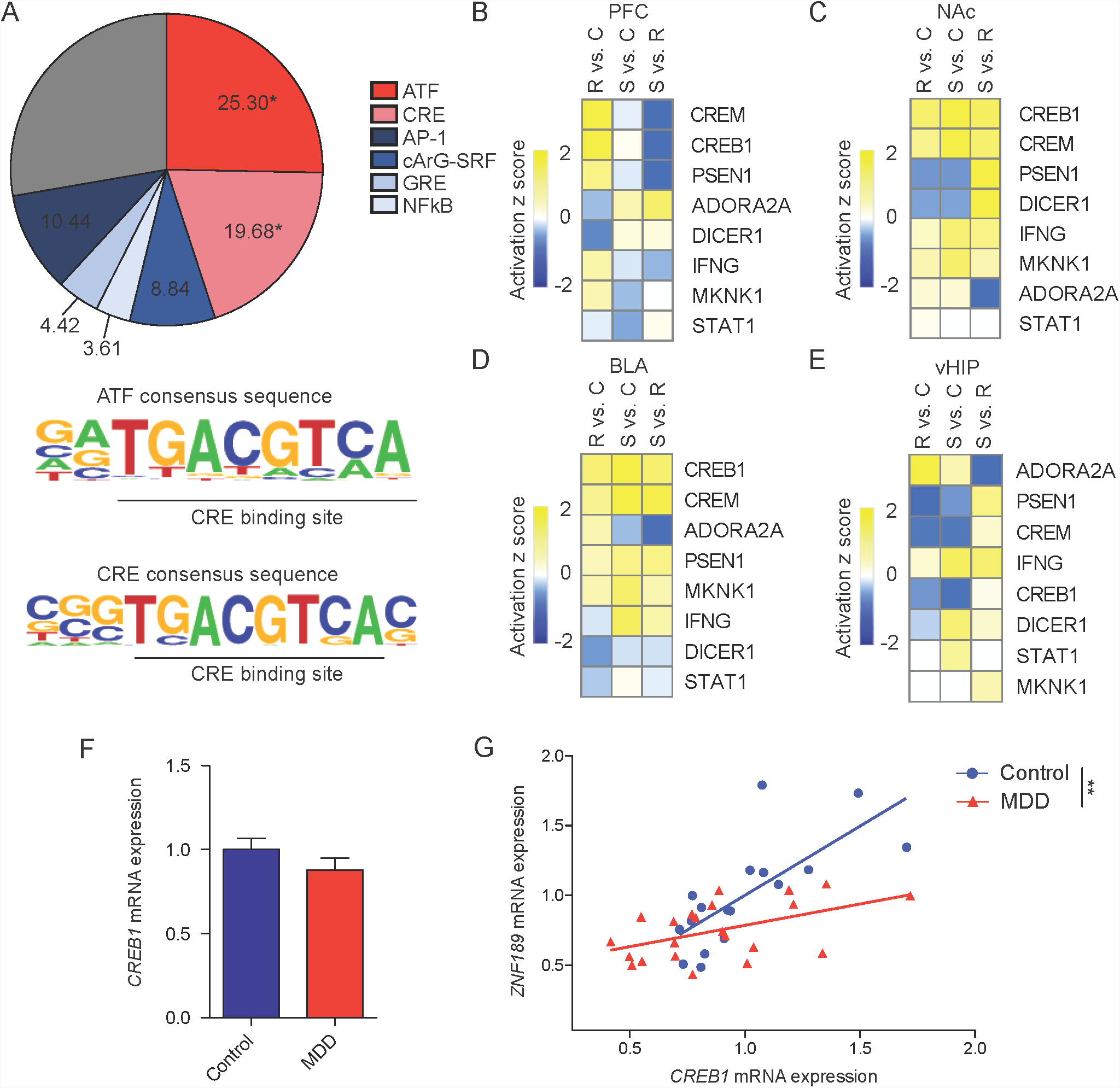
CREB is an Upstream Regulator of the Pink Module. (A) Binding motifs overrepresented (FDR q < 0.05) in the pink module (colored pink) with common non-overrepresented motifs (colored blue) included for comparison (top). Both significantly overrepresented binding motifs contain a CRE site (bottom). (B-E) Upstream regulator analysis of transcriptional changes in pink module genes. CREB is an upstream regulator across brain regions and is predicted to be upregulated in resilience in PFC. (F) mRNA levels of *CREB1* in PFC from MDD patients and matched controls (n = 17-22). (G) *CREB1* and *ZNF189* are less correlated in PFC in MDD (n = 19-22). **p < 0.01. See also Tables S2-4.

To complement this analysis, we performed an upstream regulator analysis using Ingenuity Pathways Analysis (IPA). In contrast to HOMER, which probes the promoter regions of a gene set for enrichment of known binding sites, this approach utilizes known interactions for a set of genes to determine upstream regulators based on downstream expression changes. Matching our results with HOMER, CREB is a predicted upstream regulator of pink module transcription across brain regions (Figure 3B-E; Table S4). In contrast, our HOMER finding of ATF1 regulation is not reproduced by IPA. While CREB is predicted to be upregulated more in susceptibility than resilience in nucleus accumbens (NAc), baslolateral amygdala (BLA), and ventral hippocampus (vHIP), the predicted regulation of CREB in PFC (up in resilient, no change in susceptible) mimics that of both pink module DEG enrichment and *Zfp189* differential expression (Figure 1D). As such, we hypothesized that CREB may regulate the pink module in PFC, at least in part, through interactions with *Zfp189*, which contains a CRE site in its upstream gene promoter (see below).

To investigate the role of CREB-*Zfp189* interactions in pro-resilient regulation of the pink module, we analyzed a published ChIP-chip dataset of active, Ser 133 phosphorylated CREB (pCREB) binding in NAc following CSDS (Wilkinson et al., 2009). These data reveal that pCREB binding to *Zfp189* is significantly (p < 0.001) reduced following CSDS, but administration of imipramine, a common tricyclic antidepressant, reverses this reduction (p < 0.001). To determine the relevance of these findings to PFC and human depression, we performed qPCR analysis on post-mortem PFC from the same human MDD patients and controls in which we assessed *ZNF189* (Figure 1E; Table S2). While there is no effect of MDD on *CREB1* expression (t= 1.216, p = 0.232, n = 17-22; Figure 3F), *CREB1* is related to *ZNF189* expression in both control and MDD samples (r^2^ = 0.53, p < 0.001 and r^2^ = 0.25, p = 0.018 respectively; Figure 3G), with a stronger relationship in control than MDD (F_1,35_ = 7.702, p = 0.009). As such, CREB-*Zfp189* interactions appear relevant in both human MDD and mouse CSDS.

### CREB-*Zfp189* Interactions Regulate Resilience

We next evaluated our predicted mechanism of direct CREB-*Zfp189* regulation of pink module genes and behavioral resilience. Notably, CREB manipulations in animal models of depression have been limited to NAc and hippocampus; CREB overexpression in NAc promotes depression-like behavior, while CREB knockout (KO) is pro-resilient, with opposite effects observed in hippocampus (Carlezon et al., 2005; Chen et al., 2001; Covington et al., 2011). This is consistent with our upstream regulator analysis, which predicts higher activity for CREB in NAc of susceptible mice (Figure 3C). However, our upstream regulator analysis also predicts higher CREB activity in the resilient PFC (Figure 3B); as such, CREB KO in this region should be pro-susceptible. To evaluate this prediction, we injected CREB^fl/fl^ mice with adeno-associated virus (AAV) expressing Cre recombinase plus GFP or GFP alone and exposed mice to a subthreshold social defeat procedure (Figure 4A). This defeat induced social avoidance in AAV-Cre mice but not AAV-GFP mice (F_1,26_ = 4.656, p = 0.040, n = 11-17, Bonferroni post hoc p < 0.001; Figure 4B), providing *in vivo* confirmation of our bioinformatics prediction of a pro-resilient effect of CREB in PFC. To evaluate the effects of CREB KO on *Zfp189* gene expression, we micro-dissected virally infected PFC and performed qPCR. Both *CREB1* and *Zfp189* are reduced by CREB KO (t = 2.974, p = 0.006 and U = 45.0, p = 0.036 respectively; Figure 4C-D) providing causal *in vivo* evidence of regulation of *Zfp189* by CREB in PFC.

**Figure 4.**
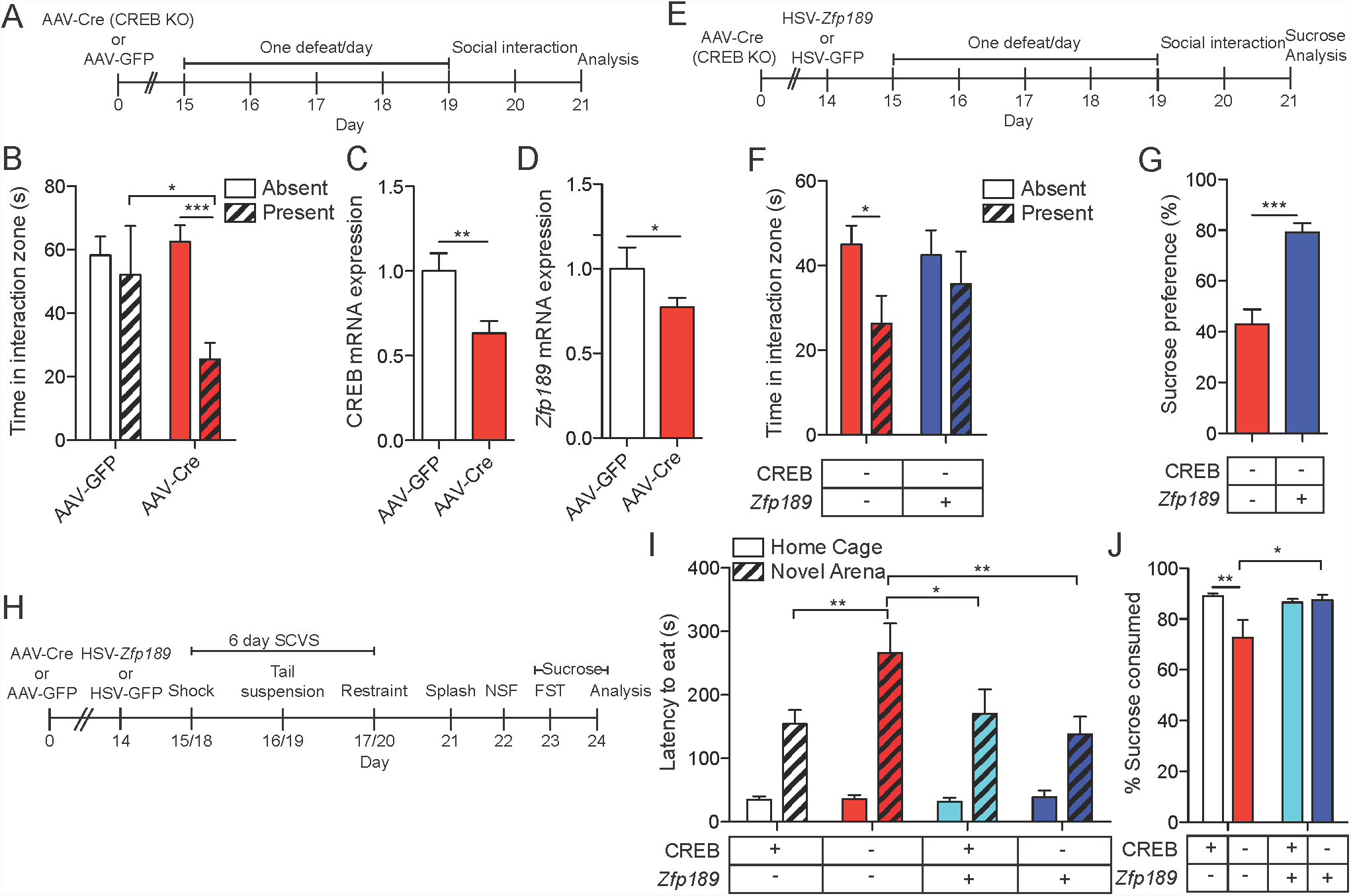
CREB Knockout (KO) in PFC Increases Susceptibility but is Rescued By *Zfp189* Overexpression. (A) Experimental timeline to evaluate behavioral effects of CREB KO in PFC. (B) Local KO of CREB in PFC produces social avoidance in SI in a subthreshold social defeat procedure (n = 11-17). (C-D) CREB KO reduces mRNA levels of (C) *Creb1* and (D) *Zfp189* (n = 11-16). (E) Experimental timeline to determine whether *Zfp189* overexpression in PFC reverses the deleterious effects of CREB KO. (F) CREB KO in PFC produces social avoidance in the SI test but concurrent overexpression of *Zfp189* reverses this deficit (n = 13-14). (G) Overexpression of *Zfp189* in PFC increases sucrose preference in CREB KO mice (n = 14). (H) Experimental schematic for female sub-chronic variable stress (SCVS) to investigate behavioral effects of CREB KO and *Zfp189* overexpression in PFC. (I) CREB KO in PFC increases latency to eat in the novel arena, but not when CREB KO is paired with *Zfp189* overexpression (n= 9-11). (J) Mice with local KO of CREB in PFC have a lower sucrose preference than control mice, an effect blocked by concurrent *Zfp189* overexpression (n = 9-11). *p < 0.05, **p < 0.01, ***p < 0.001. Bar graphs show mean ± SEM. See also Figure S4.

Since our data indicate that CREB is upstream of *Zfp189*, that *Zfp189* promotes resilience, and that *Zfp189* acts through pink module expression changes, we reasoned that CREB KO in PFC is pro-susceptible at least in part through the consequent reduction in *Zfp189*. If this is the case, concurrent overexpression of *Zfp189* should rescue the pro-susceptible effects of CREB KO. To evaluate this, we injected AAV-Cre in PFC of CREB^fl/fl^ mice while overexpressing either *Zfp189* plus GFP or GFP alone with HSV and exposed mice to a subthreshold defeat (Figure 4E). Consistent with our previous findings, CREB KO mice injected with HSV-GFP spend less time in the interaction zone when a target mouse is present (F_1,25_ = 5.690, p = 0.025, n = 13-14, Bonferroni post-hoc p < 0.05; Figure 4F). In contrast, CREB KO mice that also overexpress *Zfp189* do not exhibit social avoidance. *Zfp189* overexpressing CREB KO mice also have a higher sucrose preference than GFP overexpressing *CREB* KO mice (t = 5.176, p < 0.001, n = 13; Figure 4G), further supporting the capacity of *Zfp189* to mitigate the pro-susceptible effects of CREB KO in PFC.

Given the increasing recognition of considerable sex differences in depression, we next investigated whether CREB-*Zfp189* interactions also regulate depression-like behavior in female mice. The pink module was identified from a dataset of male mice only (Bagot et al., 2016a) and transcriptional effects of stress exhibit dramatic sex differences (Gray et al., 2015; Hodes et al., 2015; Labonte et al., 2017; LaPlant et al., 2009). Nevertheless, we found recently that another molecular regulator can induce similar behavioral effects even in the context of sex-specific transcriptional changes (Lorsch et al., 2018). This is supported clinically by observations that, despite sex-specific transcriptional differences in MDD (Labonte et al., 2017), the same antidepressants are successful in males and females (Sramek et al., 2016). Further, our identification of abnormal reduced *CREB1-ZNF189* co-expression in human MDD (Figure 3G) included both male and female samples. We injected PFC of female CREB^fl/fl^ mice with AAV-Cre plus GFP or GFP alone plus either HSV-Zfp189 or HSV-GFP and exposed mice to six days of sub-chronic variable stress (SCVS), a protocol that reliably induces depression-like behavior in female mice (Hodes et al., 2015; Labonte et al., 2017; LaPlant et al., 2009) (Figure 4H). This produced four distinct groups: AAV-GFP + HSV-GFP (CREB^+^*Zfp189*^−^), AAV-Cre + HSV-GFP (CREB^−^*Zfp189*^−^), AAV-GFP + HSV-Zfp189 (CREB^+^*Zfp189*^+^), and AAV-Cre + HSV-Zfp189 (CREB^−^ *Zfp189*^+^). In the novelty suppressed feeding (NSF) test, in which antidepressant administration reverses stress-induced increases in latency to feed in a novel environment, CREB^−^*Zfp189*^*-*^ mice show a longer latency to feed in the novel arena than CREB^−^*Zfp189*^*+*^, CREB^*+*^*Zfp189*^*-*^ and CREB^+^*Zfp189*^*+*^ mice, with no effect of CREB KO or *Zfp189* overexpression on latency to feed in the home cage (F_1,35_ = 5.301, p = 0.027, n = 9-11, Bonferroni post-hoc p < 0.01, p < 0. 01, and p < 0.05 respectively; Figure 4I). While there is no difference in grooming time in the splash test, which measures time spent grooming after application of sticky water to the fur, or latency to immobility in the FST (F_1,40_ = 0.067, p = 0.797, n = 10-12 and F_1,35_ = 0.121, p = 0.730, n = 8-11 respectively; Figure S4), CREB^−^*Zfp189*^*-*^ mice display a lower sucrose preference than either CREB^*+*^*Zfp189*^*-*^ or CREB^−^*Zfp189*^*+*^ mice (*χ*^*2*^(3) = 8.475, p = 0.037, n = 9-11, Mann-Whitney post-test p = 0.009 and p = 0.029 respectively; Figure 4J). Thus, manipulating CREB-*Zfp189* interactions in female SCVS produces similar behavioral alterations as those observed in males after CSDS.

### CRISPR-Mediated CREB-*Zfp189* Interactions Promote Resilience

Having demonstrated that *Zfp189* overexpression is sufficient to reverse the pro-susceptible effects of CREB KO in PFC, we next sought to mechanistically probe the direct interaction between CREB and *Zfp189 in vivo*. Although our data indicate a functional relationship between CREB and *Zfp189* expression (Figure 4C-D), our behavioral data could be explained by indirect interactions between the two factors. To address this question, we used CRISPR technology as a tool for gene locus-targeted recruitment of CREB to the *Zfp189* gene promoter in PFC neurons. We fused the nuclease-dead, RNA-guided, DNA-binding protein Cas9 to a constitutively active, phosphomimetic mutant form of CREB^S133D^ (dCas9-CREB^S133D^) and designed sgRNAs to target near the consensus CRE motif in the promoter region of *Zfp189*. The construct design, including the fluorescent reporters used in each vector, is shown in Figure 5A. *In vitro* validation of sgRNAs shows that the top *Zfp189-*targeting sgRNA (*Zfp189-* sgRNA) induces *Zfp189* expression only when the active CREB^S133D^, and not the phospho-null CREB^S133A^ or dCas9 with no functional moiety, is recruited to the *Zfp189* locus (U = 1.0, p = 0.009, n = 5-6; Figure S5). Our *Zfp189*-sgRNA is designed to bring dCas9-CREB^S133D^ ∼150 bp upstream of the CRE motif in the *Zfp189* promoter (Figure 5B). With the exception of the *Zfp189* locus, there is no complementary site in the mouse genome with fewer than three base mismatches, suggesting a low *in silico* probability of off-target effects (Table S5). Our control non-targeting (NT-) sgRNA is also predicted to target no specific sequence in the mouse genome (Table S6).

**Figure 5.**
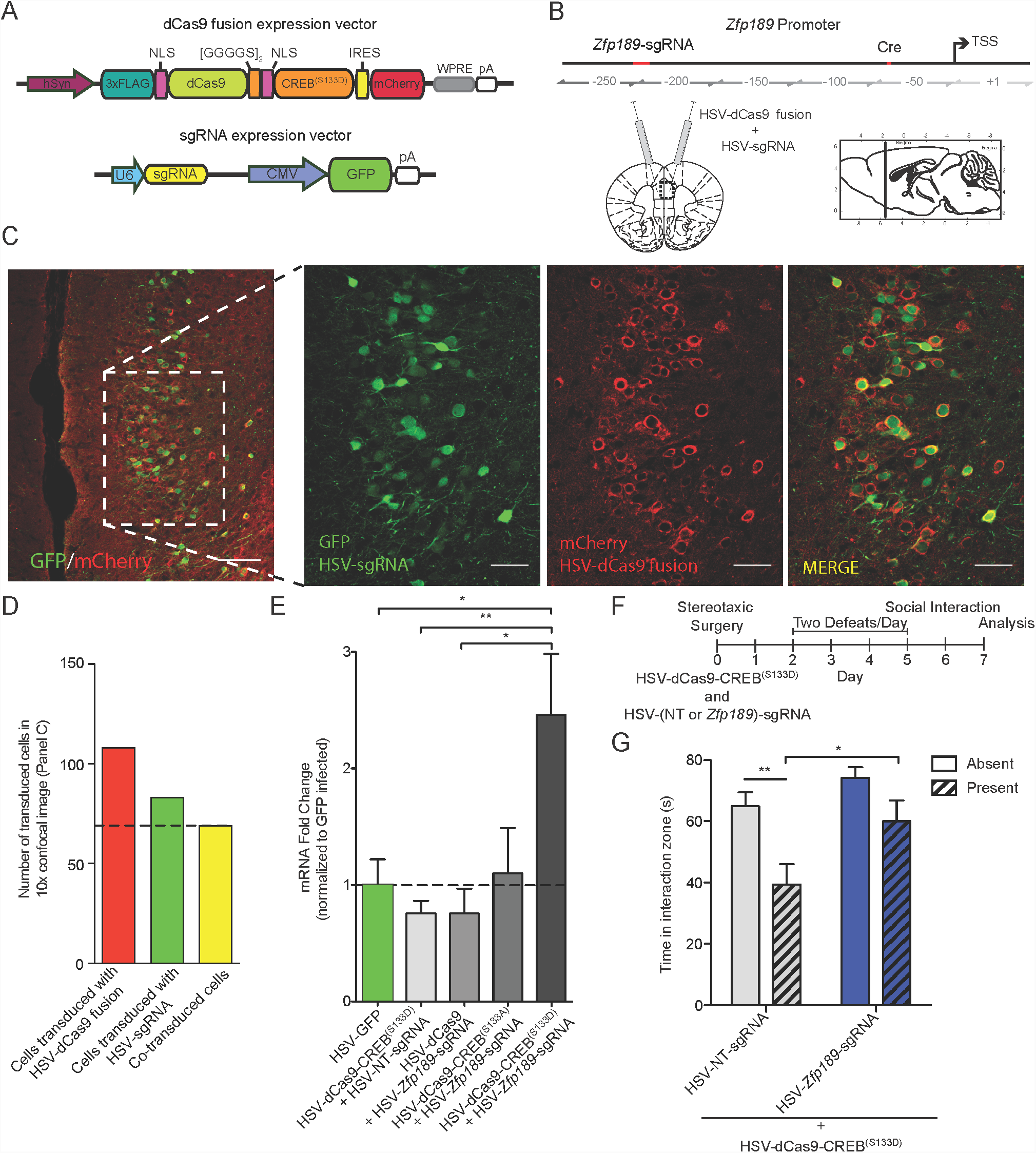
CRISPR-Mediated CREB-*Zfp189* Interactions Induce Pro-Resilient Behaviors. (A) Schematic of the CRISPR vectors. Variable dCas9 functional moiety in orange. Variable gene-targeting single guide RNA (sgRNA) in yellow. (B) Location of *Zfp189*-targeting sgRNA binding site in Zfp189 promoter. CRISPR vectors were packaged in HSV and delivered as a viral cocktail bilaterally to PFC. Hashed box denotes field of confocal imaging for Panel C, left. (C) Immunohistological staining shows high degree of colocalization of HSV-sgRNA expression vector (GFP) and HSV-dCas9 fusion expression vector (mCherry) in PFC neurons. Left: 10x objective, scale bar = 100 μm. Right: 20x objective, scale bar = 50 μm. (D) Quantification of virus colocalization. (E) CRISPR-mediated targeting of active, pseudo-phosphorylated CREB^(S133D)^ to *Zfp189* is sufficient to increase mRNA expression in PFC (n = 5-12). (F) Experimental timeline to determine effect of CRISPR-dependent CREB-*Zfp189* interactions in PFC on stress resilience. (G) Pro-resilient effects of CRISPR*-*dependent CREB-*Zfp189* interactions. dCas9-CREB^S133D^ delivered with *Zfp189-*targeting sgRNA increases time in the interaction zone when a target mouse is present relative to dCas9-CREB^S133D^ with non-targeted (NT) sgRNA (n = 38-40). *p < 0.05, **p < 0.01. Bar graphs show mean ± SEM. See also Figure S5; Tables S5-6.

We independently packaged our sgRNA expression vectors and our dCas9 fusion protein expression vectors in HSVs, and co-delivered these vectors to the mouse PFC. We observe that the two HSVs infect predominantly the same neurons, with transgene co-expression occurring in the majority of affected cells (Figure 5C-D). dCas9-CREB^S133D^ directed to the *Zfp189* promoter increases *Zfp189* expression in mouse PFC compared to HSV-GFP alone, dCas9-CREB^S133D^ paired with NT-sgRNA, and dCas9 with no functional domain paired with *Zfp189-*sgRNA (*χ*^*2*^(5) = 10.27, p = 0.036, n = 5-12; Mann-Whitney post-test p = 0.035, p = 0.004, and p = 0.040 respectively; Figure 5E). dCas9-CREB^S133A^ does not increase *Zfp189* expression, even when directed to the *Zfp189* promoter. These data demonstrate the ability to harness the physiologically-relevant mechanism of CREB-mediated induction of *Zfp189* expression with CRISPR-mediated transcriptional reprogramming in mouse brain.

We next determined whether direct CREB induction of the endogenous *Zfp189* gene locus is sufficient to increase resilience. We injected HSV-dCas9-CREB^S133D^ paired with either HSV-NT-sgRNA or HSV-*Zfp189-*sgRNA into PFC and exposed mice to an accelerated social defeat stress procedure (Figure 5F). In SI, mice delivered Cas9-CREB^S133D^ targeted to the *Zfp189* promoter spend more time in the interaction zone when a social target is present relative to mice in which Cas9-CREB^S133D^ is not targeted (NT-sgRNA) (F_1,76_ = 6.235, p = 0.015, n = 38-40, Bonferroni post-hoc p < 0.05), with only NT-sgRNA mice displaying social avoidance (Bonferroni post-hoc p < 0.01) (Figure 5G). Therefore, mimicking the endogenous interaction between pCREB and *Zfp189* in PFC is sufficient to increase resilience to social defeat.

To test our hypothesis that these effects are associated with changes in pink module gene expression, we micro-dissected virally-infected PFC and performed RNA-seq to compare the transcriptome in our defeated *Zfp189-*sgRNA mice to that of our defeated NT-sgRNA mice (Figure S6A). Our logistic regression analysis reveals activation of both the pink and green resilient modules in response to CRISPR-mediated interactions between CREB and *Zfp189* in the context of social defeat (p = 0.010 and p = 0.048 respectively; Figure 6A). While the smaller magnitude of the CRISPR effects, compared with traditional overexpression (Figure 2A), is reflected in fewer regulated pink module genes overall, these genes are predominantly upregulated, which is consistent with an activation of the pink module (Figure 6B). The smaller magnitude effects with CRISPR are likely due to the smaller magnitude of *Zfp189* induction (∼20-fold *Zfp189* induction with conventional overexpression vs. ∼2.5-fold *Zfp189* induction with CRISPR-mediated recruitment of CREB). This highlights the technical advance provided by our novel approach of mimicking a physiologically-relevant mechanism of gene induction in the brain *in vivo*.

**Figure 6.**
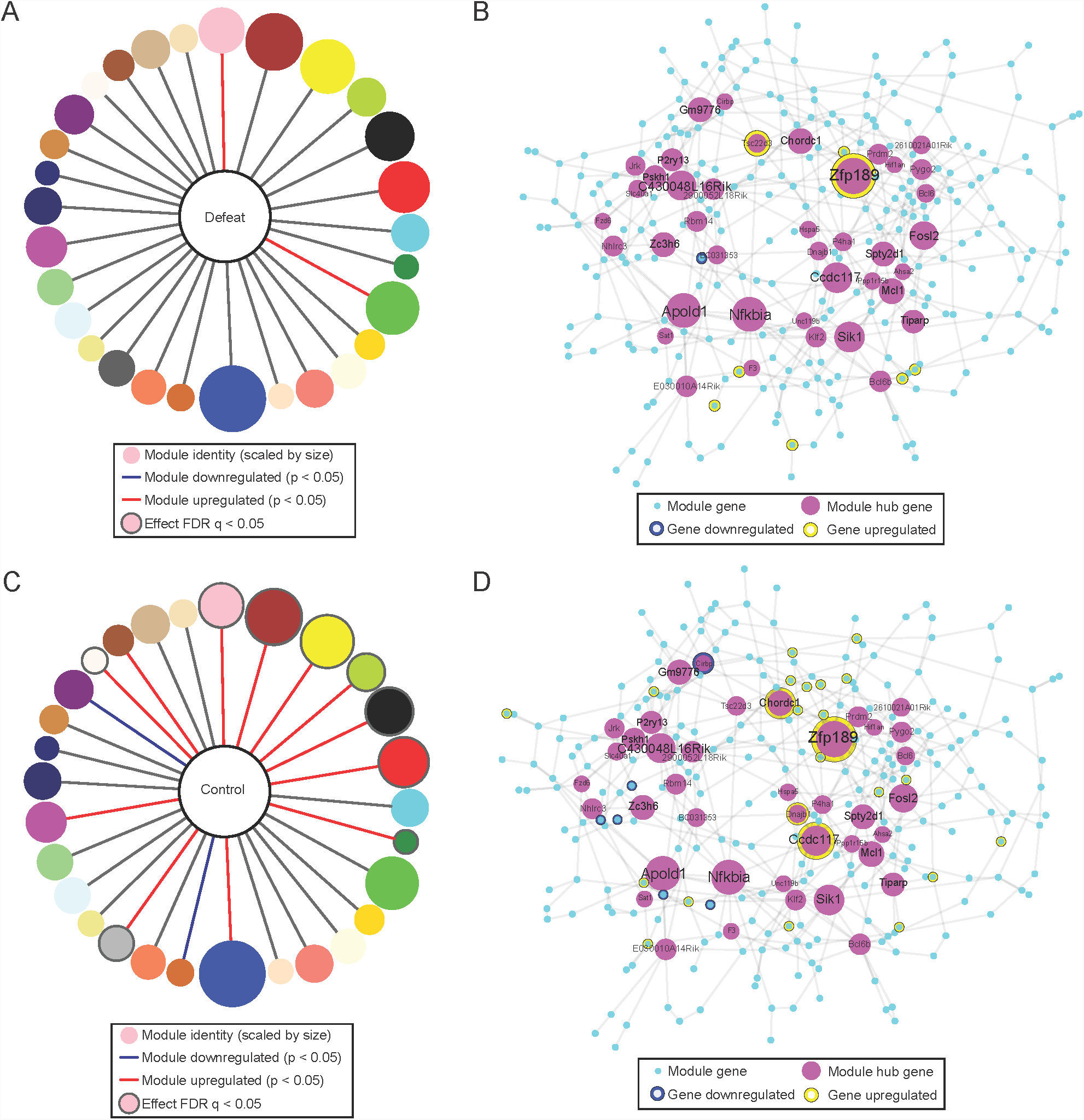
CRISPR-Induced Stress Resilience is Dependent on Pink Module Expression. (A) Module overlap for PFC DEGs (p < 0.05, log_2_FC > |0.2|) resulting from *Zfp189*-targeted dCas9-CREB^(S133D)^ compared to NT-dCas9-CREB^(S133D)^ in mice exposed to social defeat. (B) Pink module genes differentially expressed in defeated mice. (C) Overlap for PFC DEGs (p < 0.05, log_2_FC > |0.2|) resulting from *Zfp189-*targeted dCas9-CREB^S133D^ compared to NT-dCas9-CREB^S133D^ in unstressed controls. (D) Pink module genes differentially expressed in unstressed controls. See also Figure S6.

To determine whether forcing CREB-*Zfp189* interactions in PFC of unstressed controls is sufficient to produce similar changes in pink module genes, we repeated this procedure in mice not exposed to CSDS (Figure S6B). Again, the pink module is affected (p = 0.012; Figure 6C) with a predominant upregulation of pink module genes (Figure 6D). Importantly, we observe no regulation of genes nearest to the 49 most homologous off-target sites of the *Zfp189*-sgRNA across the mouse genome in unstressed control animals, and regulation of only one off target (*Tsc22d3*) in the defeat cohort, which is likely a result of the defeat experience itself (Figure S6C).

### *Zfp189* Regulates PFC Neuron Physiology

Having demonstrated that manipulations of both *Zfp189* and CREB-*Zfp189* interactions in PFC preferentially affect pink module transcription, we next sought to investigate the functional effects of these transcriptional changes. We first prepared slices from animals injected intra-PFC with HSV-*Zfp189* or HSV-GFP and recorded transduced vs. non-transduced PFC pyramidal neurons because *Zfp189* is enriched in this excitatory cell-type (Kozlenkov et al., 2016). Compared to GFP expressing neurons, neurons expressing *Zfp189* exhibit lower intrinsic membrane excitability, as they fire fewer action potentials in response to injected current (F_13,376_ = 3.186, p < 0.0001, Bonferroni post hoc p < 0.05). *Zfp189* expressing neurons also display larger amplitudes of spontaneous excitatory postsynaptic currents (sEPSCs) (F_2,38_ = 6.217, p = 0.0046, Dunnett’s post hoc p < 0.05), with no difference in sEPSC frequency (F_2,36_ = 0.2633, p = 0.7700) (Figure 7A-E). These results suggest that *Zfp189* potentiates excitatory synaptic transmission. The decreased intrinsic membrane excitability of manipulated neurons could reflect an adaptation to the *Zfp189*-mediated increase in synaptic responses, as similar intrinsic excitability homeostatic adaptations have been observed in NAc neurons (Wang et al., 2018). Since PFC pyramidal neuron activity is pro-resilient (Bagot et al., 2015; Covington et al., 2010), and stress and MDD are associated with an imbalance of excitation and inhibition in this brain region (Ghosal et al., 2017), these findings could reflect an endogenous physiological property of the resilient brain, where PFC neurons exhibit increased responses to cortical inputs.

**Figure 7.**
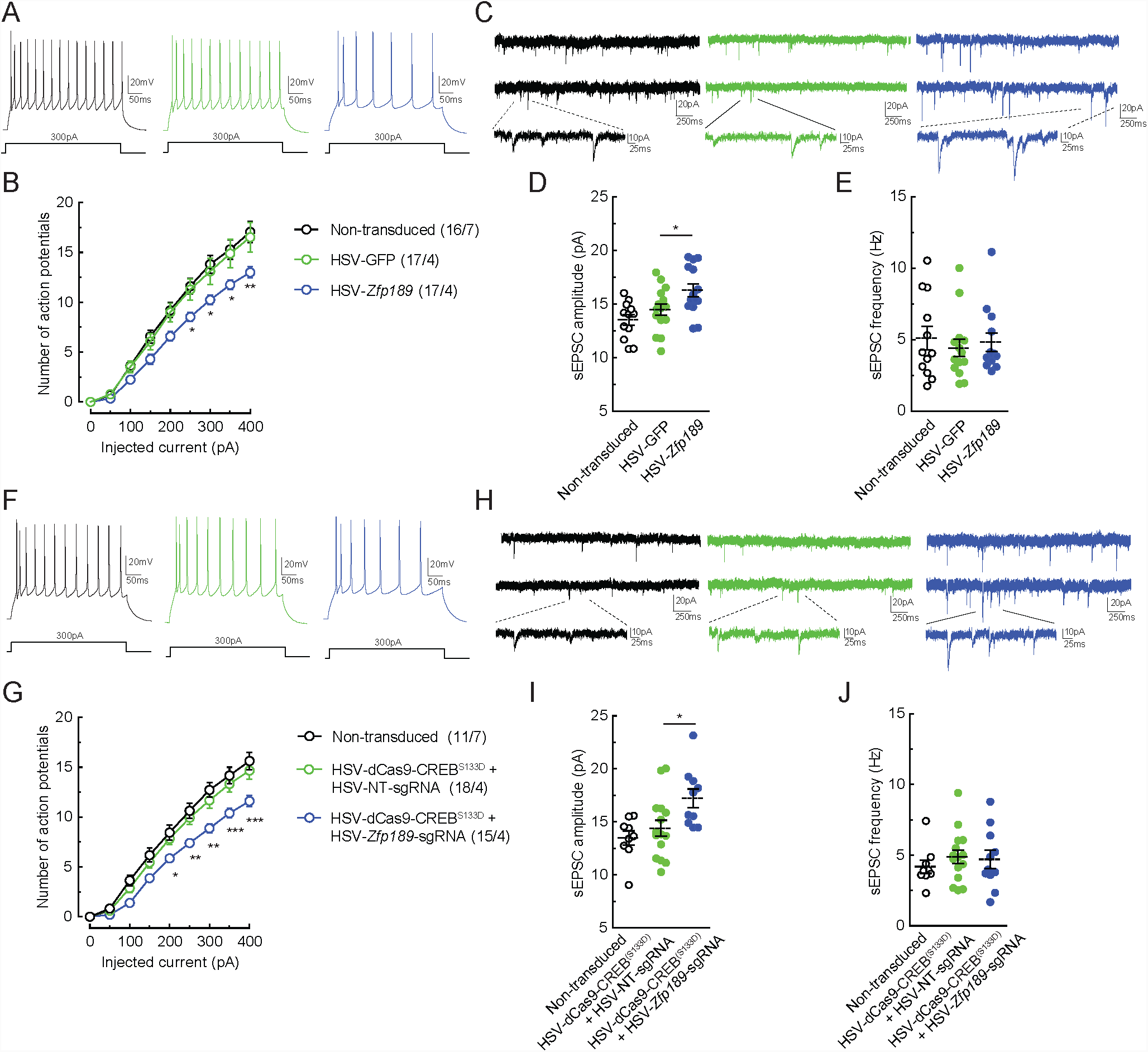
*Zfp189* and CREB-*Zfp189* Interactions Alter Physiological Properties of PFC Pyramidal Neurons. (A-E) (A) Representative recordings of (B) intrinsic excitability, and (C) representative recordings of (D-E) spontaneous excitatory postsynaptic currents (sEPSCs) in PFC pyramidal neurons infected with HSV-*Zfp189* or HSV-GFP or uninfected neurons. (F-J) (F) Representative recordings of (G) intrinsic excitability, and (H) representative recordings of (I-J) spontaneous sEPSCs in PFC pyramidal neurons infected with HSV-dCas9-CREB^S133D^ with either NT-sgRNA or *Zfp189*-sgRNA or uninfected neurons. *p < 0.05, **p < 0.01, ***p < 0.001, Dunnett’s post hoc relative to HSV-GFP or HSV-NT-sgRNA infected cells. Bar graphs show mean ± SEM.

We repeated our PFC recordings following use of our CRISPR constructs. dCas9-CREB^S133D^ paired with *Zfp189*-sgRNA similarly decreases the intrinsic membrane excitability of PFC pyramidal neurons (interaction F_16,288_ = 3.957, p < 0.0001, Bonferroni post hoc p < 0.05), with manipulated cells exhibiting larger sEPSC amplitudes (F_2,31_ = 5.302, p = 0.0105, Dunnett’s post hoc p < 0.05) and no change in sEPSC frequency compared to noninfected or GFP-expressing control cells (F_2,32_ = 0.4263, p = 0.4968) (Figure 7F-J). Thus, bringing active CREB to the *Zfp189* locus is sufficient to reproduce *Zfp189-*dependent effects on PFC pyramidal neuron physiology.

## Discussion

Here, we establish a previously unidentified pathway of stress resilience, delineating a mechanism from transcriptional network through neuronal function and behavior. We provide causal evidence that CREB binding to the *Zfp189* gene promoter induces *Zfp189* in PFC, which then activates a resilient-specific transcriptional network to ultimately mediate changes in PFC pyramidal neuron physiology and behavioral resilience. We recapitulate an endogenous interaction seen in the resilient brain with a novel CRISPR approach to provide mechanistic insight into the hierarchical regulation of gene co-expression networks and their role in regulating complex psychiatric phenotypes.

Our data indicate that while broad transcriptional changes are seen across brain regions in stress-resilient mice (Bagot et al., 2016a, 2016b; Krishnan and Nestler, 2008), complete reversal of the totality of these changes is not required to either reduce susceptibility or induce an antidepressant-like response. Manipulations of *Zfp189* influence multiple resilient networks, yet we observe that only the pink module is consistently activated across all manipulations (Figure 2B,; Figure S3B) and that variations in *Zfp189* expression affect pink module expression more than any other network (Figure 2D), suggesting that *Zfp189* preferentially exerts its resilient effects through pink module gene expression. By elucidating the regulatory hierarchy of the pink module, we show that affecting either key network genes (Figures 1-2) or vital regulatory interactions with an upstream regulator (Figure 5-6) is sufficient to affect pink module gene expression, PFC neuronal physiology, and behavioral resilience across experiments. Importantly, this is not a single-gene effect as induction of *Zfp189—*either directly or via CRISPR-mediated interactions with CREB—broadly affects downstream gene transcription. While such manipulations do not recapitulate the entire transcriptional profile of resilience, they do activate the pink module, the top ranked gene network associated with resilience (Figure 2D). These data are consistent with the view that MDD results from the coalescence of numerous, small changes in gene expression rather than the actions of single genes of strong effect (Gaiteri et al., 2014), which may explain the difficulty in finding causative variants in genome wide association studies (GWAS) in non-homogeneous populations (Ripke et al., 2013). In this regard, we do not make the claim that *Zfp189* dysregulation is the cause of MDD, but rather that *Zfp189* is uniquely situated in a network of important regulatory genes in such a way that modification of *Zfp189* is sufficient to modify a network of genes to induce a stress-resilient state.

Importantly, our identification of the pink module and *Zfp189* is both bioinformatically-driven and unbiased. The pink module is the only resilient-specific network transcriptionally active across brain regions and *Zfp189* is the top key driver gene of this network. *Zfp189* would have likely been missed by previous studies that selected genes for *in vivo* validation based on known functional pathways since *Zfp189* has not been previously implicated in any neurobiological context and little is known about the gene except for its inclusion in the Krüppel-like factor family of zinc finger proteins (Odeberg et al., 1998). The power of our approach is evidenced by data showing biological and translational relevance of the pink module, *Zfp189*, and CREB-*Zfp189* interactions. First, the pink module is enriched for PPIs and cell-type specific genes (Figure S1), increasing confidence that it is a biologically-driven functional network. Next, both *ZNF189* and the pink module are significantly reduced in human MDD (Figure 1E and Figure S1F), suggesting that loss of this network in humans is associated with depression. Finally, *CREB1-ZNF189* interactions are significantly reduced in both chronically stressed mice (Wilkinson et al., 2009) and depressed humans (Figure 3G).

In addition to defining the role of a single key driver gene within a gene network, our data explore the mechanism by which a key driver, *Zfp189*, and the associated pink module is regulated. This is an important but previously unexplored direction in the context of *in vivo* validations of network biology. Not only did we find that CREB, which has been implicated extensively in depression (Carlezon et al., 2005; Chen et al., 2001; Covington et al., 2011; Juhasz et al., 2011; Wilkinson et al., 2009; Xiao et al., 2017), is a predicted upstream regulator of the pink module with two separate analytical tools, but also that CREB enrichment is unique to the pink module. Further, we identified a previously unexplored relationship between CREB and *Zfp189*, which in animals is decreased by stress but enhanced by imipramine in NAc (Wilkinson et al., 2009) and extended these findings to PFC and human MDD (Figure 3E). This identifies a direct mechanism for pink module regulation that we explored through a series of *in vivo* manipulations: CREB bound to the *Zfp189* promoter in PFC neurons induces *Zfp189* gene expression, subsequently activating pink module genes to induce behavioral resilience. Our observation that *Zfp189* overexpression also exerts antidepressant-like effects in reversing susceptibility, and that *Zfp189* overexpression is most strongly linked to the pink module (Figure 2D), suggests that the downstream transcriptional effects of *Zfp189* are required for its antidepressant efficacy. As such, pharmacological manipulation of *Zfp189*, either directly or through CREB-dependent mechanisms, has the potential to regulate an entire network of pro-resilient genes and may be an attractive target for MDD therapeutics. Notably, the therapeutic potential for *Zfp189* is greater than for CREB itself, because CREB is involved in a wide array of cellular functions (Bleckmann et al., 2002; Shaywitz and Greenberg, 1999) and exerts pro-susceptible or pro-resilient effects in different brain regions (Carlezon et al., 2005).

Because CREB KO reduces *Zfp189* expression, and *Zfp189* can compensate for the pro-susceptible effects of CREB KO in PFC, it is likely that under physiological conditions CREB exerts its effects on the pink module in large part via *Zfp189*. Nevertheless, because we found significant CRE binding motifs on numerous pink module genes (Table S4), it is possible *Zfp189* is simply one of many pink module genes affected by CREB manipulation. Therefore, to establish the direct relationship between CREB and *Zfp189*, we used a novel CRISPR approach (Hamilton et al., 2018a). Most studies to date have relied on global overexpression or knockdown manipulations to explore the effects of a given transcription factor or epigenetic modification. However, these manipulations affect hundreds or thousands of genes simultaneously, making it impossible to determine which gene regulatory events are required for a given outcome. While our group has circumvented this problem in the past by engineering synthetic zinc finger DNA-binding proteins fused to epigenetic modifying moieties (Hamilton et al., 2018b; Heller et al., 2014, 2016), zinc-finger design can be expensive, time consuming, inflexible and difficult to predict. In contrast, CRISPR technology allows flexible and inexpensive locus-specific gene editing (Cong et al., 2013; Jinek et al., 2012; Mali et al., 2013). While CRISPR has been used previously to induce locus-specific epigenetic modifications in mammalian brain (Liu et al., 2016, 2018), our study is the first to utilize CRISPR to probe site-specific transcription factor regulation of an endogenous transcriptional network.

Our data indicate that bringing active CREB specifically to the *Zfp189* locus is not only sufficient to increase resilient behavior in mice exposed to stress (Figure 5G), but that this single molecular interaction is sufficient to activate pink module genes and affect PFC physiology (Figures 6-7). Our finding that even in the absence of stress bringing active CREB to the *Zfp189* locus in PFC neurons is sufficient to upregulate the pink module (Figure 6C) further speaks to the therapeutic potential of *Zfp189*, as the pink module, which is not present at baseline in unstressed controls (Figure 1B), can be induced by mimicking the CREB-*Zfp189* interactions observed in resilient mice. These data represent an important technical advance in available approaches, detail the regulatory role of CREB and signify the first time CRISPR has been used *in vivo* in brain to deliver a transcription factor to a single gene locus to manipulate behavior. Moreover, these data provide a further understanding of the mechanisms involved in network biology by illustrating that modulating a single in-network molecular regulator, or an upstream regulator that interacts with that gene, is sufficient to fundamentally alter expression of a larger gene network. While the role of CREB in the context of MDD is not entirely explained through its interactions with *Zfp189*, our evidence showing a causal relationship in the context of stress resilience indicates that CREB-*Zfp189* interactions in PFC are important for a resilient state. Still, further studies are needed to dissect the *Zfp189-*dependent effects of CREB from its broader profile of molecular actions.

In sum, we describe a vital single molecular interaction of the known transcriptional regulator CREB and the novel downstream transcription factor *Zfp189* that is capable of activating a network of genes in PFC to mediate stress resilience. These findings elucidate the molecular mechanisms involved in stress resilience, as well as provide a broad molecular framework for the hierarchical organization and regulation of gene co-expression networks and their relationship to complex behavior.

## Acknowledgments

This work was supported by National Institutes of Health grants F30MH110073 (Z.S.L.), T32GM007280 (Z.S.L.), T32MH096678 (Z.S.L.), K99DA045795 (P.J.H), P50MH096890 and R01MH051399 (E.J.N.) and the Hope for Depression Research Foundation.

## Author Contributions

Conceptualization, Z.S.L, P.J.H, R.C.B, and E.J.N; Methodology, Z.S.L, P.J.H, I. O. T, S.M, I.M, and E.J.N; Software, Z.S.L, A.R, A.M, X.Z, Y.E.L, B.Z, and L.S; Formal Analysis, Z.S.L, P.J.H, A.R, X.Z, Y.E.L, B.Z, and L.S; Investigation: Z.S.L, P.J.H, E.M.P, M.S, A.L, P.M, O.I, L.A, S.P, B.L, A.C, and A.E.S; Resources, A.M, R.N, and G.T; Writing – Original Draft, Z.S.L and P.J.H; Writing – Review & Editing, Z.S.L, P.J.H, R.C.B., and E.J.N; Visualization, Z.S.L, P.J.H, M.S., and X.Z; Supervision, I.M, B.Z, L.S, R.C.B, and E.J.N; Project Administration and Research Funding, E.J.N.

## Declaration of Interests

All authors declare no competing financial interests.

## Contact For Reagent and Resource Sharing

Further information and requests for resources and reagents should be directed to and will be fulfilled by the Lead Contact, Eric Nestler

## Experimental Model and Subject Details

### Animals

Experimental mice were either C57BL/6J mice or C57BL/6J mice genetically modified for a conditional brain region-specific Cre-dependent CREB knockout by insertion of *loxP* sites flanking *Creb1* exon 2 (Covington et al., 2011). In addition, 6-month-old CD1 aggressor mice were used as aggressors to induce social stress in the chronic social defeat stress (CSDS) procedure, but were not included in any analysis. Wildtype C57BL/6J mice were 8 weeks old at the time of experimentation. CREB^fl/fl^ mice were used between 8 weeks and 4 months of age due to breeding considerations, and age was counterbalanced across experimental conditions. C57BL/6J mice were housed 5 per cage; CD1 mice were single-housed. C57BL/6J mice undergoing CSDS were single-housed following the final defeat and C57BL/6J mice undergoing SCVS were single-housed following the final stressor. In order to maintain consistent study design, unstressed controls were single-housed at the same time as stressed mice. Once mice were single-housed, they remained as such until the end of experimentation. For all experiments, mice were randomized to experimental groups and only healthy, well-appearing mice were selected for experimentation. All mice were maintained on a 12-hour light-dark cycle with lights on at 7:00 AM and a controlled temperature range of 22-25°C. Food and water were provided *ad libitum* except for the 24 hours preceding novelty suppressed feeding (NSF) testing, when food was removed. All experiments conformed to the Institutional Animal Care and Use Committee (IACUC) guidelines at Mount Sinai. Behavioral testing took place during the animals’ light cycle. In cases where non-automated analysis was used, experimenter was blinded to experimental group. Order of testing in behavioral experiments was counterbalanced and assignment to experimental groups was random.

## Method Details

### Stress Protocols and Behavioral Testing

CSDS and social interaction tests were performed according to established protocols (Berton et al., 2006; Krishnan et al., 2007). CD1 retired breeder mice were screened for aggression in three-minute intervals over the course of three days. CD1 mice consistently attacking 8-week-old male C57BL/6J screener mice were included as aggressors for CSDS. On the first day of stress, CD1 mice were placed on one side of a large hamster cage separated by a perforated Plexiglas divider and an 8-week-old male C57BL/6J was placed on the other side. Importantly, this Plexiglas divider allows for sensory, but not physical, contact between CD1 and C57BL/6J mice. During each defeat, C57BL/6J mice were placed in the same side of the cage as the CD1 aggressor for a period of 7.5-10 minutes. This duration was kept constant throughout each experiment and was predetermined to titrate the defeat based on the overall aggression in the CD1 cohort during screening. During this time, the CD1 aggressor physically attacked the C57BL/6J mouse. Following the physical bout, C57BL/6J mice were returned to the other (empty) side of the divider where they remained in sensory contact with the CD1 aggressor that had just attacked them, but could not be further harmed physically. For each consecutive defeat session (24 hours later for CSDS and subthreshold defeat and 12 hours later for accelerated defeat), the C57BL/6J mouse was exposed to a new CD1 aggressor in a different hamster cage and the procedure was repeated. Control C57BL/6J mouse were double-housed in a mouse cage separated by a perforated divider for the length of stress. To control for handling effects, control mice were moved to the adjacent half cage whenever a stress occurred for experimental mice. CSDS took place for a duration of 10 days, accelerated defeat took place over the course of 4 days with 2 defeats per day (to coincide with the time course of HSV-mediated transgene expression), and subthreshold defeat took place over the course of 5 days.

SCVS was performed as previously described (Hodes et al., 2015) with three different hour-long stresses performed twice over a total of six days. Briefly, 8-week-old female C57BL/6J mice were exposed to foot shock (0.45 mA) on days one and four, tail suspension on days two and five, and restraint stress (in a 50 ml Falcon tube in the home cage) on days three and six. Mice were group housed (five mice/cage) when they were not being stressed and control mice remained in their home cages throughout.

Behavioral tests occurred in a behavior suite different from where stress exposure was performed. In cases where a test was repeated following a manipulation, a different behavior room was used for the second test. Mice were given one hour to habituate to the behavioral room prior to behavioral testing. Due to the timeline of HSV expression (Neve et al., 2005), multiple behaviors occurred on the same day when HSV vectors were used. In order to minimize spillover effects from one test to another, tests were separated by a minimum interval of 2 hours. Behavioral analysis for social interaction, locomotion and open field test (OFT) was performed automatically by video tracking software (Ethovision 10.0, Noldus), FST was analyzed manually on pre-recorded video by investigators blind to study design and sucrose preference test and NSF were analyzed manually in real time. To ensure adequate power, sample sizes were chosen in accordance with number of mice needed to show statistical significance in CSDS and SCVS as defined by previous studies (Bagot et al., 2016a; Hodes et al., 2015).

Social interaction testing was performed under red light 24 hours after the last social defeat stress. C57BL/6J mice were placed into an open arena with an empty wire cage at one side (interaction zone). Mice were given 2.5 minutes to explore the arena and then removed. A novel CD1 aggressor to which the C57BL/6J mouse had never been exposed to was placed in the cage (interaction zone) and the procedure was repeated. Time in the interaction zone was recorded automatically with video tracking software. Data were analyzed as time spent in the interaction zone when the aggressor was absent compared to time spent in the interaction zone when the aggressor was present. In cases where mice were subset into resilient and susceptible phenotypes, defeated mice with social interaction ratio scores > 1.2 that spent > 60 seconds in the interaction zone when the target was present were determined to be resilient. Defeated mice with SI ratio scores < 0.8 that spent < 40 seconds in the interaction zone when the target was present were determined to be susceptible.

OFT was performed by allowing mice 10 minutes to explore an open arena under red light. Although nothing was physically placed in the arena, center and periphery were defined in video tracking software. Total time in center was recorded and utilized for analysis. In addition, total distance moved during this time was analyzed to determine locomotor effects.

Forced swim test (FST) was always performed last in the sequence of behavior. FST was performed in 4L Pyrex beakers filled with two liters of 25**°**C (± 1°) water. Mice were placed into the water and recorded by a front-facing camera for a period of six minutes. Investigators blind to study design scored FST videos by recording latency to the first immobility state.

NSF was performed following 24 hours of food deprivation. Female mice were placed in a novel arena with corncob bedding and a single piece of food in the center of the arena. Time to feed was recorded manually under white light. Mice were given a maximum of 10 minutes to eat, after which the trial was ended and latency of 600 seconds was recorded. After the mouse ate in the novel arena, the mouse was returned to the home cage where a single piece of food was located in the center, and time to eat in the home cage was recorded. Data were analyzed as latency to eat in the novel arena and latency to eat in the home cage.

Sucrose preference test was performed as a two-bottle choice test. One bottle was filled with water and the other bottle was filled with 2% sucrose. Initial weights of each bottle were recorded and bottle weights were recorded each morning and evening over the sucrose preference period. Sucrose preference was calculated as change in weight of the sucrose bottle/change in weight of both bottles X 100. Total sucrose preference was used for analysis.

In order to allow for peak viral expression at the time of behavior, adeno-associated virus (AAV) infusions and behavior were separated by four weeks and herpes simplex virus (HSV) viral infusions and behavior were separated by 4-5 days as described in reported experimental timelines.

### Tissue Collection

Mice were euthanized 24 hours after final behavior via cervical dislocation. In order to acquire only infected tissue within prefrontal cortex (PFC), microdissection was performed under a fluorescent microscope. Brain slices containing PFC were either visualized on blade and punched directly or suspended in cold phosphate buffered saline (PBS) prior to tissue dissection. PFC was collected as either a single midline 12-gauge punch or bilateral 14-gauge punches, which varied according to virus spread. However, only PFC was included in dissections. Mice were excluded from analysis when the PFC was not correctly targeted. Dissected tissue was immediately frozen on dry ice. Since our HSV and AAV vectors express green fluorescent protein (GFP), there was no way to distinguish expression of the two viruses. As such, for experiments in which HSV and AAV vectors were both injected, PFC was collected with a 12-gauge punch and downstream quantitative reverse-transcription polymerase chain reaction (qPCR) was used to validate virus effects. In cases where dual 14-gauge punches were used, two punches (bilateral) from each mouse were combined, but samples were never pooled between mice.

### Viral Reagents

We overexpressed *Zfp189* using bicistronic p1005 HSV expressing GFP alone or GFP plus *Zfp189*. This involves a dual promoter approach whereby GFP expression is driven by a cytomegalovirus (CMV) promoter but *Zfp189* expression is driven by IE4/5. *Zfp189* was inserted into the p1005 plasmid from a plasmid containing the mouse *Zfp189* gene (Origene MR209370), which was packaged into HSV.

We overexpressed Cre recombinase in CREB^fl/fl^ mice using AAV serotype 2 AAV-CMV-Cre-GFP virus from the University of North Carolina Vector core. Similar to expression of *Zfp189*, this virus induces Cre expression via the CMV promoter. In CREB^fl/fl^ mice where we intended to preserve CREB expression, we injected AAV-CMV-GFP (serotype 2).

We repurposed the CRISPR system in order to target CREB binding to the *Zfp189* promoter. We cloned and synthesized a novel construct of a phosphomimetic mutant form of CREB fused to nuclease-dead *Staphylococcus pyogenes* Cas9 protein (dCas9-CREB^S133D^), which can localize to specific sites along the genome based on the complementarity of a specific single guide RNA (sgRNA) sequence. sgRNA sequences for *Zfp189* were determined first *in silica* based on the sequence of the *Zfp189* promotor and a suite of 10 sgRNAs were designed to bind to distinct DNA sequences proximal to the CRE motif within the promoter. To promote specificity, sgRNA off-target effects were predicted at crispr.mit.edu according to their published algorithm (Hsu et al., 2013), and candidate sgRNAs with complementary sequences in the mouse genome with fewer than three base mismatches were excluded. The sequence of our non-targeting sgRNA (protospacer sequence: GCGAGGTATTCGGCTCCGCG) was identified from the GeCKO v2 libraries and validated to ensure that no genomic site was targeted. All 10 candidate sgRNAs were *de novo* synthesized as gBlocks from Integrated DNA Technologies containing a U6 promoter, variable target sequence, guide RNA scaffold and termination signal. These were sub-cloned into a p1005 variant plasmid for HSV packaging. All *Zfp189-*sgRNAs were first validated in N2A cells to identify the most effective *Zfp189*-targeting sgRNA. Cells were transfected and lysed after 48 hours and mRNA expression was analyzed with qPCR. The top sgRNA (protospacer sequence: GTGTCTCGGTTAGCAAGAAG) was packaged into HSV and tested *in vivo*.

*In vivo* confirmation of *Zfp189* induction was validated via qPCR on dissected PFC tissue. In order to minimize between-mouse variability, a hemispheric approach was utilized wherein the test constructs were injected into one hemisphere while control constructs was injected into the other hemisphere of the same mouse.

### Viral-Mediated Gene Transfer

Stereotaxic surgeries targeting PFC were performed as previously described (Bagot et al., 2016a; Hamilton et al., 2018c). Mice were anesthetized with an intraperitoneal injection of ketamine (100 mg/kg) and xylazine (10 mg/kg) dissolved in sterile water. Subsequently, mice were placed in a small-animal stereotaxic device (Kopf Instruments) and the skull surface was exposed. 33-gauge needles (Hamilton) were utilized to infuse 0.5 µL of virus at a rate of 0.1 µL/minute followed by a five-minute rest period to prevent backflow. For CREB^fl/fl^ and CRISPR experiments, 1 µL of virus at a rate of 0.2 µL/minute was utilized to maximize viral spread within the PFC. The following coordinates were utilized for PFC (from Bregma: anterior/posterior +1.8 mm, medial/lateral +0.75 mm, dorsal/ventral −2.7 mm; 15° angle). While PFC injections targeted the infralimbic cortex, virus spread sometimes extended beyond these anatomical boundaries to other PFC regions.

### Immunohistochemistry

Mice were anesthetized with an intraperitoneal injection of ketamine (100 mg/kg) and xylazine (10 mg/kg) and transcardially perfused with a fixative solution containing 4% PFA paraformaldehyde (PFA) (w/v) in 0.1 M Na_2_HPO_4_/ Na_2_HPO_4_, pH 7.5 at 4°C delivered at 20 ml/min for 5 min with a peristaltic pump. Brains were post-fixed for 24 hours in 4% PFA at 4°C. Sections of 30 µm thickness were cut in the frontal plane with a vibratome (Leica, Nussloch, Germany) and stored at −20°C in a solution containing 30% ethylene glycol (v/v), 30% glycerol (v/v) and 0.1 M phosphate buffer. Free-floating sections were processed for immunohistochemistry as follows. On day one, sections were rinsed three times for 10 min in PBS before permeabilization for 15 min in PBS containing 0.2% Triton X-100 (Fisher). Sections were then rinsed three times in PBS followed by a blocking step of one hour incubation in PBS containing 3% bovine serum albumin. Primary antibodies against GFP (Avès Lab, GFP-1020, 1:500 dilution), mCherry (Abcam, ab125096, 1:500 dilution) were diluted in blocking solution and sections incubated overnight at 4°C with gentle shaking. Sections were then washed three times in PBS and incubated with secondary antibodies (donkey anti-chicken Alexa Fluor 488, donkey anti-mouse Cy3; Jackson ImmunoResearch, 1:500 dilution) for two hours at room temperature. After three rinses in PBS, sections were mounted in Vectashield (Vector Labs).

GFP and mCherry expression was assessed in PFC using a LSM 710 laser-scanning confocal microscope (Carl Zeiss) imaged using a 10X or 20X objective with a 1.0 digital zoom.

### RNA Isolation, qPCR, Library preparation, and Sequencing

Total RNA was isolated from frozen dissected PFC tissue using QIAzol lysis reagent and purified using the miRNAeasy mini kit (Qiagen). Following isolation, RNA to be utilized in qPCR was quantified by Nano Drop (Thermo Fisher) and converted to cDNA with iScript (Bio-Rad). qPCR samples were analyzed in triplicate using the standard ΔΔCT method. For non-CRISPR experiments, hypoxanthine phosphoribosyltransferase 1 (*Hprt1*) was utilized for normalization for both mice and humans, and for CRISPR experiments, gene expression was normalized to the geometric mean of *Hprt1*, dCas9, and GFP transcripts to account for both cell health and appropriate delivery of tool components. For RNA being utilized for RNA-seq, RNA integrity (RIN) was assayed using an Aligent 2100 Bioanalyzer (Aligent, Santa Clara CA). Average RIN values were above nine and samples with RIN values less than eight were excluded from analysis. Each sample consisted of PFC punches from the same animal with no pooling between animals. RNA sequencing of the effects of *Zfp189* expression included five samples per group (five HSV-*Zfp189* and five HSV-GFP) for previously susceptible mice and 4-5 samples per group (five HSV-*Zfp189* and four HSV-GFP) for unstressed controls. RNA sequencing of CRISPR-infected tissue included three samples per group (three dCas9-CREB^(S133D)^ + NT-sgRNA and three dCas9-CREB^(S133D)^ + *Zfp189*-sgRNA) for undefeated controls and 8-12 samples per group (12 dCas9-CREB^(S133D)^ + NT-sgRNA and eight dCas9-CREB^(S133D)^ + *Zfp189*-sgRNA) for defeated mice. Libraries were prepared using the TruSeq RNA Sample Prep Kit v2 (Illumina, San Diego CA). Briefly, mRNA was polyA selected from the total RNA pool. mRNA was then fragmented and converted to cDNA with reverse transcriptase followed by cDNA size selection and purification with AMPure XP beads (Beckman Coulter, Brea CA). To identify each sample, strand-specific adapters were ligated to adenylated 3’ ends and an additional size selection step was performed. The cDNA library was then amplified using polymerase chain reaction. During this step, barcodes of six base pairs were added to the adaptors. Library quality and concentration was measured by the Bioanalyzer before sequencing. Libraries were sequenced by either the Genomics Core Facility of the Icahn School of Medicine at Mount Sinai (*Zfp189* overexpression) using an Illumina HiSeq 2500 System with v3 chemistry and 100 base pair single-end reads or GENEWIZ (CRISPR sequencing) using a Illumina HiSeq System with 150 base pair paired-end reads. Multiplexing was performed to ensure minimum reads of 20 million for each sample.

### RNA-seq

Raw reads obtained from PFC were mapped to mm10 using HISAT2 (Kim et al., 2015). SAM files were converted to BAM files and were sorted according to chromosome number using SAMTools (Li et al., 2009). Counts of reads mapped to genes were obtained using HTSeq-count (Anders et al., 2015) against Ensembl v90 annotation. Differential expression analysis was carried out using the R package DESeq2 (Love et al., 2014). For virally infected tissue, sequenced samples in which the overexpressed transgene could not be adequately detected (*Zfp189* for HSV-*Zfp189* or dCas9, sgRNA, or *Zfp189* for CRISPR studies) were considered to be a result of experimenter error (viral targeting or tissue selection) and were removed from analysis.

### Quantification and Statistical Analysis

#### Identification of Resilient-Specific Co-Expression Networks

In order to identify gene networks implicated in resilience, we utilized a previously published dataset from our group that reported weighted gene co-expression network analysis (WGCNA) modules and differentially expressed genes following CSDS (Bagot et al., 2016a). Briefly, WGCNA modules and differential expression profiles were generated from RNA-seq data with tissue taken at 3 time-points after 10 days of CSDS. Mice were exposed to stress, phenotyped in the social interaction test, and the PFC, nucleus accumbens (NAc), ventral hippocampus (vHIP), and basolateral amygdala (BLA) were dissected to for RNA-seq. Differential expression comparisons were region-specific but brain regions were pooled prior to WGCNA. As such, reported WGCNA networks represent relationships of genes across brain regions.

Our approach to identifying resilient-specific co-expression networks was predicated on our hypothesis that resilient-specific modules would be both unique to the resilient phenotype and transcriptionally active immediately following CSDS. As such, we first examined the 30 resilient modules for module differential connectivity (MDC) (Zhang et al., 2013). MDC is a measure of how connectedness amongst a set of genes is altered in the same genes in a different condition. This was performed as previously described (Bagot et al., 2016a) and modules with FDR q < 0.05 are reported as significantly differentially connected. However, while previous studies from our group (Bagot et al., 2016a; Labonte et al., 2017) identified networks for functional validation based on MDC in a single condition, gene expression (Bagot et al., 2016a; Wilkinson et al., 2009) and circuit (Friedman et al., 2014) alterations in the resilient phenotype are distinct from that of susceptibility or control and therefore, only networks that showed MDC when compared to both phenotypes were considered resilient-specific.

### Module Enrichment Analysis

In order to identify statistical overlap between previously identified differentially expressed genes (Gene Expression Omnibus database: GSE72343) (Bagot et al., 2016a) and modules, we utilized the Super Exact Test (SET) R package (Wang et al., 2015). The SET evaluates multi-set interactions to determine the difference between observed and expected overlap, which is then quantified statistically as an enrichment p-value and fold change. Expected overlap for two gene sets is dependent on the size of the gene sets and the number of total variables in the dataset (background number of genes). Higher than expected overlaps are indicated by larger fold change values, which represents the ratio of observed overlap to expected overlap. SET is advantageous in this setting as enrichment for upregulated and downregulated genes can be determined separately. Similarly, we used SET to evaluate enrichment for cell-type specific genes. Cell-type specific genes were determined based on a meta analysis of multiple public RNAseq datasets, each available at www.celltypes.org. The top 200 genes for each cell-type were used in enrichment testing. Protein-protein interactions were determined using STRING version 10.5 (Szklarczyk et al., 2017).

### Human Brain Analysis

To identify whether our resilient mouse modules were preserved in human brain, we used the modulePreservation function of R package WGCNA to compare module identity in mouse to RNA sequencing data from human control and human MDD (GEO dataset: GSE102556) (Labonte et al., 2017). All brain regions included in the human analysis were included, and male and female subjects were combined. To evaluate the difference between preservation in human controls and human MDD, we compared module preservation p-values in each condition. A 10-fold change in p-value was considered to be a detectable deviation.

To evaluate *CREB1* and *ZNF189* in human brain, we used qPCR on reverse-transcribed mRNA from BA25, the PFC region most homologus to the ventral medial PFC targeted in our mouse studies. Samples from individuals with alcohol in their blood at the time of death were excluded due to possible effects on the transcriptome in PFC. Demographics of final cohort are shown in Table S2. There were no significant differences in any demographic between MDD and control.

### Identification of the Pink Module Network Structure

We generated a network structure for the pink module using the Algorithm for the Reconstruction of Accurate Cellular Networks (ARACNE) (Margolin et al., 2006). Critically, however, ARACNE is not a directed analysis, so connections between genes are still based on correlations. In order to attempt to resolve regulatory genes within the pink module, we next performed Key Driver Analysis (KDA) (Zhang and Zhu, 2013) on the ARACNE reconstructed network. KDA is predicated on the understanding that more important regulatory genes will have a larger effect on other genes in the module. Therefore, more important genes should forge more direct connections in the final module structure than less important genes. We analyzed the pink module structure at a threshold of 2 layers to identify the most important regulatory genes. Key driver genes had a number of connections significantly above the average value for the network and were considered for further *in vivo* analysis.

### Identification of the Relationship Between Zfp189 and the Pink Module

To establish the relationship between Zfp189 and the WGCNA modules, we first obtained gene expression from DESeq2’s regularized log-transformation. Each gene’s expression was then standardized to have zero mean and unit standard deviation across all samples. PCA was performed on each module (excluding Zfp189) to obtain the first principal component after weighing as the “eigen-gene” for each module. The Pearson’s correlation analysis was then performed between Zfp189 and each module’s eigen-gene to get the R^2^ score.

### Determination of Pink Module Upstream Regulators

To probe the pink module for specific regulatory binding sites, we utilized HOMER motif analysis (Heinz et al., 2010). HOMER examines binding sites within a gene set to see if a specific binding motif is significantly enriched compared to what would be expected in a background gene set (in our case, the entire mouse genome). HOMER motif analysis was performed on all resilient modules. As such, we utilized a FDR cutoff of p < 0.05 for significant upstream regulators. To reduce the detection of false positives, we limited our candidates to known binding motifs.

To complement motif analysis, we performed upstream regulator analysis using the upstream regulator tool in QIAGEN’s Ingenuity^®^ Pathway Analysis (QIAGEN Redwood City, www.qiagen.com/ingenuity) (IPA). This function predicts the identity and direction of change of known upstream regulators for a given differential expression signature from the magnitude and scale of gene expression changes in a dataset. Predictions used in this study were based on experimentally observed interactions within all datasets in IPA with the stringent filter setting applied. Reported p-value calculations were determined from the Ingenuity Knowledge Base reference set considering both direct and indirect relationships. Any regulator for which there was sufficient evidence to generate an activation/inhibition prediction is reported. Input to IPA was prepared from transcriptional changes determined by RNA sequencing in 48 hours after CSDS (Gene Expression Omnibus database: GSE72343) (Bagot et al., 2016a). Data were filtered for protein-coding genes. Three separate comparisons (susceptible vs. resilient, resilient vs. control, and susceptible vs. control) were utilized and fold change values for all pink module genes were included as input.

### Evaluation of Overlap Between RNA-seq and Resilient Modules

To test the overlap between RNA-seq differential list and the resilient modules, we used multinomial logistic regression to predict the module membership using gene expression changes as regressors. The gene expression changes are log fold changes (LFCs) from the differential analysis. To control for noise, we combined p-values with LFCs by forcing them to zero when the p-value is less than 0.05 to derive a so-called p-value adjusted LFC (PLFC). To control for covariates, such as gene length, GC contenct, etc., that may affect the differential analysis, we included the log-transformed basal gene expression (LBGE) as a covariate in the logistic regression. The LBGE is standardized to have zero mean and the same standard deviation as PLFC to facilitate optimization. In multinomial logistic regression, we used the turquoise module as the reference since it is the largest module and does not show overlap with the differential list. The coefficient of the PLFC from the regression analysis can be interpreted as the significance of the overlap while the coefficient of the LBGE indicates the bias of the covariates.

### GeneOntology

Gene Ontology for Biological Pathways (GOBP) was determined in EnrichR with gene identities of differentially expressed genes (Chen et al., 2013).

#### Preparation of PFC Slices and Electrophysiology Recordings

12-24 hours after viral infusion, mice were deeply anesthetized with isofluorane and decapitated. Coronal slices (250 µm) containing the PFC were prepared on a VT1200S vibratome (Leica, Germany) in 4°C cutting solution containing (in mM) 135 N-methyl-D-glutamine, 1 KCl, 1.2 KH_2_PO_4_, 0.5 CaCl_2_, 1.5 MgCl_2_, 20 choline-HCO_3_, and 11 glucose, saturated with 95% O_2_/5% CO_2_, pH 7.4 (305-310mM mOsm). Slices were incubated in artificial cerebrospinal fluid (aCSF) containing (in mM) 119 NaCl, 2.5 KCl, 2.5 CaCl_2_, 1.3 MgCl_2_, 1 NaH_2_PO_4_, 26.2 NaHCO_3_, and 11 glucose, saturated with 95% O_2_/5% CO_2_ (285-290mM mOsm), for 30min at 37°C. Slices in aCSF were then allowed to recover for 30 min at room temperature before electrophysiological recordings.

All recordings were made from pyramidal neurons with large, tear-dropped-shaped somas from the ventral medial PFC. Interneurons, with small round somas, were avoided and excluded from data collection. In addition to morphological criteria, action potential firing pattern was also used to differentiate between pyramidal neurons and interneurons (Yang et al., 1996). Neurons were visually targeted using DIC/fluorescence microscopy (Olympus, BX-51), with virally transfected neurons identified by GFP fluorescence (HSV-*Zfp189*, HSV-GFP) and the presence of GFP and mCherry (CRISPR). Whole-cell patch clamp recordings were made using a Multiclamp 700B amplifier and Digidata 1440A digitizer (Molecular Devices) through boroscilicate glass electrodes (2-5MΩ). Recordings were filtered at 3 kHz, amplified 5 times, and digitized at 20 kHz using Clampfit 10.2 (Molecular Devices). Series resistance was 8-25 MΩ, uncompensated, and monitored throughout the recordings. Cells with a change in series resistance >15% were excluded for analysis.

During recordings, slices were superfused with aCSF, heated to 29-31°C by passing solution through a feedback controlled in-line heater (Warner, CT). 100 µm picrotoxin was included in the bath aCSF to isolate excitatory postsynaptic currents (EPSCs). To measure intrinsic excitability, whole-cell patch clamp recordings were obtained under voltage clamp at −80 mV with a potassium-based internal solution (in mM: 130 K-methanesulfate, 10 KCl, 10 Hepes, 0.4 EGTA, 2 MgCl_2_, 3 Mg-ATP, 0.25 Na-GTP; pH 7.3, 290mM mOsm). Following a period of stabilization (∼3-5min), recordings were switched to current-clamp configuration with resting membrane potential adjusted to ∼80mV by injecting a small current. A current step protocol (from −200 to +400pA, with 50pA increments; 400ms pulse duration, 10s interpulse interval) was then repeated at least 3 times to ensure stable evoked action potential firing. After returning to voltage-clamp mode, neurons were recorded in gap-free mode to detect spontaneous EPSCs (sEPSCs) for 3 minutes. sEPSC amplitude and frequency were analyzed with Minianalysis Program (Synaptosoft Inc.).

### Statistics

Statistics were performed in Prism version 5.0 for Mac (GraphPad Software, La Jolla CA) and SPSS Statistics version 22 (IBM Corp, Armonk NY). For all behavioral analyses, outlier detection was performed using a Grubbs test with an alpha value of 0.05 and statistical outliers were excluded from analysis. Because there is no non-parametric equivalent for repeated measures tests, in accordance with the literature all SI and NSF data were analyzed using a mixed model analysis of variance (ANOVA) where one of the factors (i.e., target present, target absent) was treated as a repeated measure. A three-factor analysis model was employed when there were either two behavioral conditions (control vs. stress or pre-test vs. post-test) or two viruses (HSV and AAV) in addition to the repeated measure. In other cases, a two-factor analysis model was employed. For all tests other than SI and NSF, Levene’s test of variance was first utilized to ensure that the data met the assumptions necessary for parametric statistics. In cases where data met the assumptions necessary for parametric statistics, data from two groups were analyzed using a two-tailed students t-test and data from three or more groups were analyzed with a one-way or two-way ANOVA. In cases where the data did not meet assumptions for parametric statistics, data from two groups were analyzed using an independent samples Mann-Whitney and data from three or more groups were analyzed with a Kruskal-Wallis test. In these cases different groups were compared by independent samples Mann-Whitney tests. For parametric statistics, Bonferroni tests were used as post-tests. For electrophysiology, a Dunnett’s post-test was used instead to compare all groups to the transfected (HSV-GFP or HSV-NT-sgRNA). For comparisons of two continuous variables, we used linear regression analysis, in which case the coefficient of determination (R^2^) is reported. To compare linear regressions, we used analysis of covariance (ANCOVA).

### Data and Software Availability

RNA-seq data reported in the paper is deposited in GEO with the accession number: PENDING.

## Supplemental Information titles and legends

**Figure S1. Gene Module Biological Validity and Disease-Relevance, Related to Figure 1**

(A) Resilient module enrichment for known and predicted protein-protein interactions (PPIs). Presence of color indicates statistical significance (FDR q < 0.05) with intensity of color scaled according to -log_10_(p-value).

(B) Resilient module enrichment for the top 200 cell-type specific genes generated from a meta-analysis of cell-type specific datasets. Presence of color indicates statistical significance (p < 0.05) with intensity of color scaled according to -log_10_(p-value). Asterisk denotes FDR q < 0.05.

(C-D) Module preservation analysis in RNA-seq data from post-mortem brain from (C) human controls and (D) MDD patients. Dotted line represents significance threshold (Bonferroni corrected p < 0.05).

(E) Resilient module preservation in human controls and MDD. Presence of color indicates statistical significance (Bonferroni p < 0.05) with intensity of color scaled according to -log_10_(p-value).

(F) Change in resilient module preservation between human controls and MDD. Modules showing more preservation in controls (difference between control and MDD - log 10 p-value > 1) are shown in blue. Modules showing more preservation in MDD (difference between control and MDD -log 10 p-value < −1) are shown in red. Modules not significantly (Bonferroni corrected p < 0.05) preserved or preserved similarly (difference in |-log 10 (p-value)| < 1) are not assigned a color.

**Figure 2. Pink Module Resilient Characteristics and *Zfp189* Behavior, Related to Figure 1**

(A) Pink module genes overlaid with genes upregulated (yellow) and downregulated (blue) in the differentially expressed gene (DEG) comparison resilient vs. control in PFC (p < 0.05, FC > 1.3).

(A) Connections between pink module key driver genes.

(C) Correspondence between differential expression for the top 10 key driver genes and DEG enrichment for the pink module across phenotypes and brain areas. Presence of color indicates statistical significance (p q < 0.05) with intensity of color scaled according to -log_10_(p-value).

(D) Injection of HSV-*Zfp189* in PFC increases *Zfp189* mRNA relative to HSV-GFP (t=2.718, p = 0.024, n = 10).

(E) Exposure to CSDS reduces OFT exploration in HSV-GFP mice, but not HSV-*Zfp189* mice. However, unstressed mice show baseline differences (n = 8-14).

(F) Defeat reduces locomotion regardless of whether *Zfp189* is overexpressed (n = 9-14).

(G) Null effects of *Zfp189* overexpression in FST (n = 9-14).

(H-J) Null effects in (H) OFT, (I) locomotor behavior, and (J) FST) for previously susceptible mice injected with *Zfp189* in PFC (n = 12-13).

*p < 0.05, **p < 0.01, ***p < 0.001. Bar graphs show mean ± SEM.

**Figure S3. *Zfp189* Overexpression Upregulates the Pink Module in the Absence of Stress, Related to Figure 2**

(A) DEGs (p < 0.05, FC > log_2_|0.2|) in PFC in *Zfp189* overexpressing unstressed controls.

(B) Module-wide enrichment for HSV-*Zfp189* overexpression in PFC in unstressed mice.

(C-D) Gene Ontology Biological Process (GOBP) pathways affected *Zfp189* overexpression in PFC in (C) previously susceptible mice and (D) unstressed controls (DEG threshold = p < 0.05, FC > log_2_|0.2|).

*FDR q < 0.05

**Figure S4. CREB-*Zfp189* Effects in Splash Test and FST in Females, Related to Figure 4**

(A) Effect of CREB knockout (KO) and *Zfp189* overexpression in NSF (n = 10-12).

(B) Effect of CREB KO and *Zfp189* overexpression in FST (n = 8-11).

**Figure S5. CRISPR-mediated Overexpression of *Zfp189* in N2A Cells, Related to Figure 5**

(A) Targeting dCas9 with no functional domain to the *Zfp189* promoter does not affect *Zfp189* expression (U = 4.0, p = 0.343, n = 6).

(B) Targeting dCas9 fused to phosphomimetic CREB^S133D^ increases *Zfp189* (U = 1.0, p = 0.009, n = 5-6).

(C) Localizing dCas9 fused to a phospho-null CREB^S133A^ does not affect *Zfp189* expression (t=0.9868, p = 0.362, n = 4).

**Figure S6. Transcriptional Effects of CRISPR CREB-*Zfp189* Interactions in PFC, Related to Figure 6**

(A-B) GOBP pathways affected by dCas9-CREB^S133D^ combined with *Zfp189-*sgRNA in PFC in (A) mice exposed to social defeat and (B) unstressed controls. (DEG threshold = p < 0.05, FC > log_2_|0.2|, Asterisk = FDR q < 0.05)

(C) On and off-target CRISPR gene regulation in RNA-seq datasets. Expression of the targeted *Zfp189* gene is significantly up-regulated (p < 0.05, FC > log_2_|0.2|) in both defeated and un-stressed mice. The other 49 genes analyzed were those in closest proximity to other genomic regions with closest homology to the *Zfp189-*sgRNA used. No genomic region contains fewer than 3 mismatches with this sgRNA. Only one of these predicted off-target sites is affected by dCas9-CREB^S133D^ plus *Zfp189-*sgRNA, but this regulation was only seen in defeated mice, suggesting that this is not an off-target effect of the CRISPR tools, but rather an effect of the defeat per se.

**Table S1, Related to Figure 1. Pink Module Genes and Key Drivers.**

List of pink module genes and status as a key driver from KDA (key driver analysis). Key driver genes are ranked by number of connections two layers deep in network structure, which is included.

**Table S2, Related to Figures 1 and 3. Human Cohort Demographics.**

Gender, ethnicity, history of abuse, cause of death, age, postmortem interval (PMI), brain pH, and presence of antidepressants, alcohol, and drugs of abuse in blood of human samples used in analysis.

**Table S3, Related to Figure 3. Upstream Motifs of Module Genes.**

HOMER-defined upstream regulators for genes in the pink, blue, gold, red, and turquoise modules, which are the only modules with significant (FDR q < 0.05) upstream regulators. Included are the probed DNA sequence, number and percent of in-module genes containing sequence, number and percent of background genes containing sequence, and significance of overlap.

**Table S4, Related to Figure 3. IPA Associations of CREB and Pink Module Genes.** IPA-annotated associations between CREB and genes within the pink module used to designate CREB as an upstream regulator.

**Table S5, Related to Figure 5, *Zfp189-*Targeting sgRNA Predicted Binding Sites Throughout *Mus Musculus* Genome.**

Except for at the targeted *Zfp189* locus, there are no high-affinity binding sites for our *Zfp189*-targeting sgRNA across the entire mouse genome. Of the 141 predicted off-targets, the top sites with “scores”>0.3 are represented here. “Score” ranges from 0-100 and denotes the predicted on-target faithfulness of our synthesized sgRNA acting at the mismatched locus. The closest “matches” exhibit >3 mismatched nucleotides.

**Table S6, Related to Figure 5, Non-Targeting sgRNA Protospacer Predicted Binding Sites Throughout *Mus Musculus* Genome.**

There are no high-affinity binding sites for our NT-sgRNA across the entire mouse genome. All of the 15 total low-probability sites and “scores” are presented above. “Score” ranges from 0-100 and denotes the predicted on-target faithfulness of our synthesized sgRNA acting at the mismatched locus.

